# Engaging Biological Oscillators through Second Messenger Pathways Permits Emergence of a Robust Gastric Slow-Wave during Peristalsis

**DOI:** 10.1101/2021.06.19.449120

**Authors:** Md Ashfaq Ahmed, Sharmila Venugopal, Ranu Jung

## Abstract

Peristalsis, the coordinated contraction - relaxation of the muscles of the stomach is important for normal gastric motility and is impaired in motility disorders. Coordinated electrical depolarizations that originate and propagate within a network of interconnected layers of interstitial cells of Cajal (ICC) and smooth muscle (SM) cells of the stomach wall as a slow-wave, underly peristalsis. Normally, the gastric slow-wave oscillates with a single period and uniform rostrocaudal lag, exhibiting network entrainment. Understanding of the integrative role of neurotransmission and intercellular coupling in the propagation of an entrained gastric slow-wave, important for understanding motility disorders, however, remains incomplete. Using a computational framework constituted of a novel gastric motility network (GMN) model we address the hypothesis that engaging biological oscillators (i.e., ICCs) by constitutive gap junction coupling mechanisms and enteric neural stimulus activated signals can confer a robust entrained gastric slow-wave. We demonstrate that while a decreasing enteric neural stimulus gradient that modulates the intracellular IP_3_ concentration in the ICCs can guide the aboral slow-wave propagation essential for peristalsis, engaging ICCs by recruiting the exchange of second messengers (inositol trisphosphate (IP_3_) and Ca^2+^) ensures a robust entrained longitudinal slow-wave, even in the presence of biological variability in coupling strengths. Our GMN with the distinct intercellular coupling in conjunction with the intracellular feedback pathways and a rostrocaudal enteric neural stimulus gradient allows gastric slow waves to oscillate with a moderate range of frequencies and to propagate with a broad range of velocities, thus preventing decoupling observed in motility disorders. Overall, the findings provide a mechanistic explanation for the emergence of decoupled slow waves associated with motility impairments of the stomach, offer directions for future experiments and theoretical work, and can potentially aid in the design of new interventional pharmacological and neuromodulation device treatments for addressing gastric motility disorders.

**Author Summary:** The coordinated contraction and relaxation of the muscles of the stomach, known as peristalsis is important for normal gastric motility and primarily governed by electrical depolarizations that originate and propagate within a network of interconnected layers of interstitial cells of Cajal (ICCs) and smooth muscle cells of the stomach wall as a slow-wave. Under normal conditions, a gastric slow-wave oscillates with a single period and uniform rostrocaudal lag, exhibiting network entrainment. However, the understanding of intrinsic and extrinsic mechanisms that ensure propagation of a robust entrained slow-wave remains incomplete. Here, using a computational framework, we show that in conjunction with an enteric neural stimulus gradient along the rostrocaudal ICC chain, and intercellular electrical coupling, the intercellular exchange of inositol trisphosphate between ICCs prevents decoupling by extending the longitudinal entrainment range along the stomach wall, even when variability in intercellular coupling exists. The findings from our study indicate ways that ensure the rostrocaudal spread of a robust gastric slow-wave and provide a mechanistic explanation for the emergence of decoupled slow waves associated with motility impairments of the stomach.

## Introduction

Gastric peristalsis, the coordinated contraction and relaxation of the muscles of the stomach, is a critical phenomenon for food propulsion and waste product elimination [1] and is impaired in motility disorders [2–4]. The coordination of the contractions along the rostrocaudal compartments of the stomach causes rhythmic longitudinally travelling aboral muscle contractions [5]. Sometimes in motility disorders, the contractions can occur at a faster rate than the usual (tachygastria) or at a slower rate (bradygastria) [4,6]; whereas in some cases the rostrocaudal coordination of contractions is lost resulting in decoupling (the activity of the caudal end and the rostral end become independent of each other) or functional uncoupling (the activity of the caudal end controlling the activity of the rostral end [7,8]) of contractions in the stomach [9]. Events like these have been associated with gastric motility disorders such as gastroparesis (delay in food transit), functional dyspepsia and gastroesophageal reflux disease [4,6,10].

Peristalsis is governed by diverse and overlapping mechanisms which normally ensure robust motility patterns. Peristalsis emerges from a mutually coupled chain of pacemaker cells called the interstitial cells of Cajal (ICCs). The ICCs can independently generate electrical activity known as pacemaker potentials, which drive rhythmic potentials in the circular and longitudinal smooth muscle (SM) cells embedded in the stomach wall [5,11–13]. The electrical activity propagates through this network of cells in a coordinated manner resulting in gastric slow-wave (GSW) propagation [13,14] that underlies peristalsis.

The propagation is enabled in part by gap junction channels between the ICCs, between ICCs and SM cells, as well as between SM cells [15–17]. The SM cells, lacking intrinsic pacemaker capability, do not regenerate the slow-wave. Regeneration of the gastric slow-wave instead occurs within the pacemaker ICCs. Without coordinated ICC-ICC interactions, gastric slow-wave events do not propagate long distances [18]. Therefore, understanding how coordination arises between ICCs for propagating the gastric slow-wave is critical for understanding abnormal peristalsis that is present in gastric motility disorders.

A causal relationship between electrical gap-junction coupling in a network of ICCs and the emergent gastric slow-wave has been unequivocally demonstrated by computational studies (e.g., [7,8,19]). Pacemaker ICCs contain calcium (Ca^2+^) and inositol 1,4,5-trisphosphate (IP_3_) within their cytoplasm [20,21]. Gap junctions also facilitate exchange of second messengers such as Ca^2+^ and IP_3_ between adjacent cells [22–25]. Such exchange of Ca^2+^ and IP_3_ can impact intracellular concentration of Ca^2+^ and IP_3_ within pacemaker cells and therefore, can modulate the oscillation frequency of the concerned cell. In addition, the intracellular concentration of IP_3_ is modulated by the enteric neural stimuli received by the ICC [26–29]. Therefore, understanding the role of second messengers and their neural control is likely important for understanding aberrations in slow-wave propagation that result in the emergence of gastric motility disorders.

The longitudinally arranged ICCs along the stomach wall with intercellular gap junctions, resemble a chain of coupled oscillators [30,31]. In such a chain, the rostral or leading oscillator can gradually engage the trailing oscillators along the chain such that, at a steady-state, all the oscillators are frequency and phase-locked, with a constant phase lag between consecutive oscillators [32–34]. This phenomenon known as oscillator entrainment is thought to be the basis for the GSW in the stomach [30,31]. The integrative role of neurotransmission and intercellular coupling mechanisms in GSW entrainment remain incompletely understood. To distinguish the intercellular exchange of second messengers from passive electrical conductance is empirically challenging due to lack of pharmacological or genetic tools. The second messengers, including Ca^2+^ and IP_3_ that can move passively via gap junctions [22,23,25,35,36], are involved in intercellular communication via a spatiotemporal spread of coordinated oscillations in gap-junction coupled networks [22,23,25,35–37] and can increase the GSW frequency [38]. Yet, it is not clear whether their intercellular gap junction permeabilities can enable regenerative movement from the leading to the trailing ICCs to entrain cells and thus facilitate propagation of the slow-wave [8,18].

Our objectives were therefore to first develop a computational framework which includes constitutive stomach wall cells, biophysical models of the cells, gap junctions through which electrical coupling exists, second messenger exchange across gap junctions, and modulation of second messenger concentration by an endogenous enteric neural stimulus. Second, we utilized this framework to assess the contribution of intercellular electrical coupling and intercellular exchange of second messengers on longitudinal entrainment of the gastric slow-wave.

To develop a computational framework, we created a gastric motility network (GMN). We modeled realistic cell models for ICCs incorporating the intrinsic mechanisms for pacemaker potential generation (or oscillatory) activity. To enable inter-ICC coordination, we modeled electrical coupling and second messenger (IP_3_ and Ca^2+^) exchange between adjacent cells in a chain. Each ICC was also coupled to an SM cell, forming a pacemaker unit. The membrane voltage and period (or frequency) of the SM cells were used to assess network entrainment, as in empirical studies. Further, we set a rostrocaudal gradient for the enteric neural stimulus input to the ICCs along the ICC chain.

Since enteric neural stimulation influences the intracellular second messenger concentration, we first evaluated the importance of a gradient in the stimulus along the chain for development of a caudally propagating gastric slow-wave. We hypothesized that a linear gradient would enable gastric slow-wave propagation along the chain. Next, we examined whether exchange of second messengers alone can generate propagation of the gastric slow-wave. We hypothesized that addition of exchange of second messengers to electrical coupling would enhance the length of the oscillator chain over which entrainment is preserved (referred to as *entrainment range* hereafter). When the oscillators spanning the rostral end of the chain to the caudal end of the chain are entrained, we refer to the phenomena as rostrocaudal entrainment and if the entrained region spans a certain length from the rostral end, but falls short of the caudal end, we considered it partial entrainment. We also assessed how changing electrical coupling strength and second messenger permeability influenced the pacemaker potential frequency and the time for the slow-wave to propagate from the rostral to the caudal end of the stomach, i.e. the velocity of the slow-wave. We hypothesized that systematic increase in coupling strengths would both increase the pacemaker potential frequency and reduce the time taken for gastric slow-wave propagation. Finally, we hypothesized that variability in coupling strengths from cell-to-cell would not disrupt longitudinal entrainment if both electrical coupling and second messenger exchange mechanisms are present.

A novel computational framework with both pacemaker and muscle cells that includes intercellular exchange of second messengers and where the pacemakers receive a neural stimulus was developed to simulate and understand propagation of the slow-wave in the stomach. The results indicate that a gradient of neural modulation along the ICCs is necessary for gastric slow-wave propagation and its presence controls the directionality of the propagating slow-wave in an entrained network. Although intercellular exchange of second messengers is not necessary for slow-wave propagation, its presence can enhance the rostrocaudal length of the stomach over which entrainment is preserved and, in its absence, the entrainment range is compromised (partial entrainment is observed). This compromise is reflected by signs of decoupled slow waves and bradygastria. As hypothesized, on an increase of electrical coupling strength and second messenger permeability, the velocity of slow-wave propagation increased while the pacemaker potential frequency increased with the former and decreased with the latter. Our model with the distinct intercellular mechanisms (exchange of second messengers and electrical coupling) in combination with the intracellular feedback pathways and a rostrocaudal neural stimulus gradient allows SM cells to oscillate with a moderate range of frequencies and the gastric slow-wave to propagate with a broad range of velocities. Importantly, in the presence of variability in coupling strengths as would occur in biological networks, the existence of intercellular exchange of IP_3_ can preserve the longitudinal entrainment to a greater extent along the length of the stomach and eliminate signs of bradygastria and/or tachygastria. Together these results enhance our understanding of the intrinsic and extrinsic mechanisms engaging second messengers for the propagation of a robust gastric slow-wave essential for normal peristalsis.

## Results

### A biologically realistic gastric motility network (GMN) model for the stomach

To develop a gastric motility network (GMN) model for the stomach, we considered the length of the stomach spanning the mid-corpus to the terminal antrum. The schematic of the stomach in **Fig 1A** highlights the arrangement and diversity of cell-types in the stomach wall [11,12]. We focused on modeling the myenteric ICC and circular muscle layers which are most widely studied experimentally. The computational models for the ICCs and SM cells include a diverse set of biophysical properties reported in these cells [39–41] and are formulated as conductance-based models [42,43]. In the pacemaker ICCs, we incorporated widely accepted pathways for producing intrinsic oscillatory behavior. These included a cytosolic mitochondrial-endoplasmic reticulum (ER)-based Ca^2+^ buffering mechanism, a membrane potential dependent intracellular concentration change in IP_3_, and IP_3_ receptor mediated Ca^2+^ release from the ER, in addition to the transmembrane voltage-activated ionic currents (see **Equations 3 and 4** in **Methods**) as shown in **Fig 1B** [20,41]. Whereas for the SM cell model, we included a muscle-specific sarcoplasmic-reticular mechanism of Na^+^/Ca^2+^ exchange to moderate the intracellular Ca^2+^ in addition to the transmembrane currents as noted in **Fig 1C**. The ionic current equations defining the properties of these cells were derived from published models for the ICCs and SM cells (e.g., [42,43]) and rely on relevant experimental work for the validity of the underlying model assumptions (e.g., [39,40,44]). These cellular features were incorporated in our model to ensure both biological compliance as well as agreement with published computational models (e.g., [7,19,42,43], see **S1 Table**). Our equations, parameter descriptions, and simulation methods are described in the **Methods** section. The resulting simulated membrane voltage characteristics bear close resemblance to empirical evidence, as shown in **Fig 1D (left**, ICC pacemaker potential**; right**, SM cell slow-wave potential). The intrinsic frequency of oscillations for both ICCs and SM cells were tuned to ∼3.0 cycles per minute to match closely with the membrane potential recordings from the guinea-pig gastric antrum ICCs [5] and the canine antrum SM cells [40].

**Fig 1.**
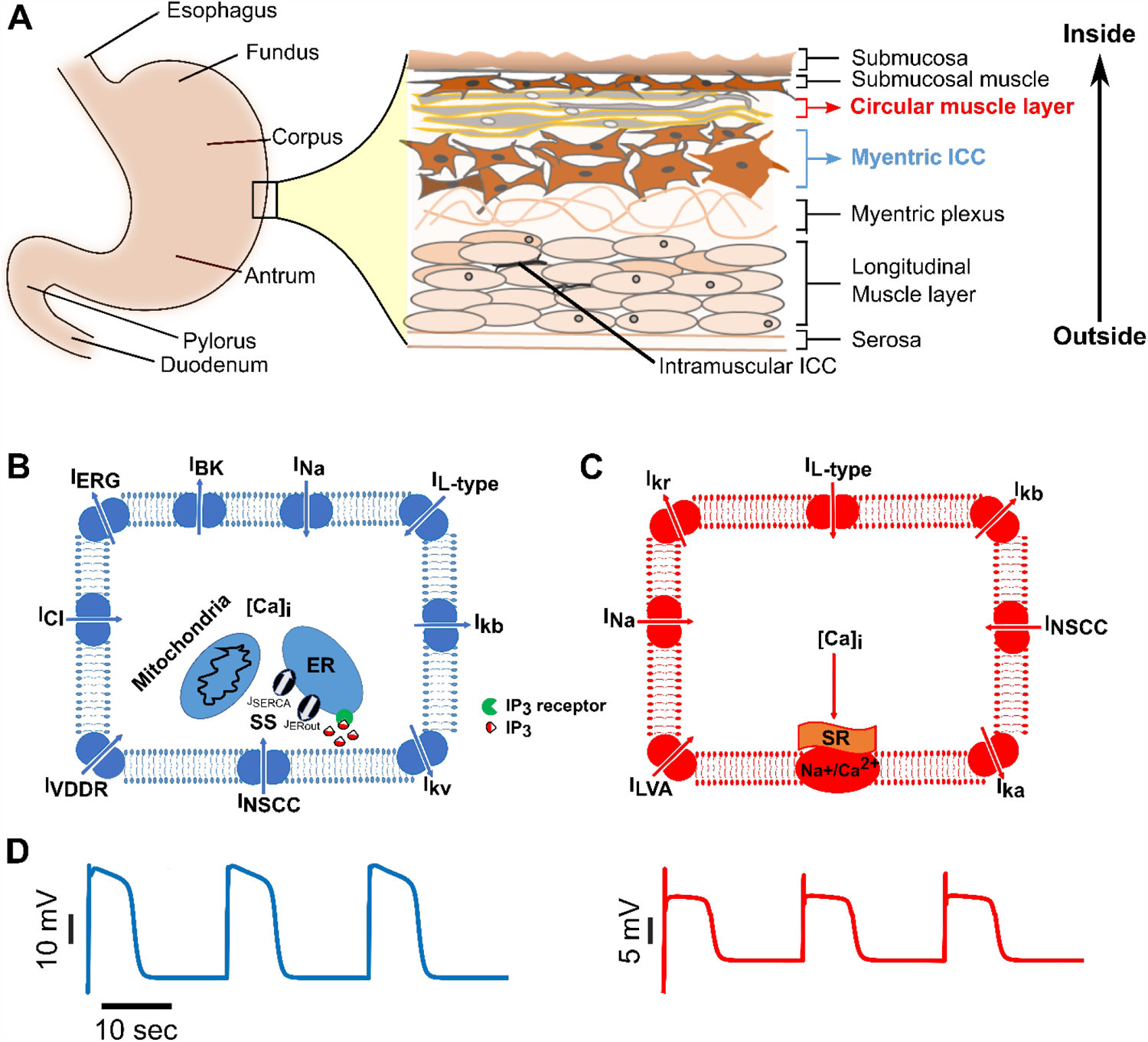
Biophysical models for ICCs and SM cells involved in gastric motility. Schematics showing **(A)** anatomical divisions of the stomach and the arrangement of various cell types in the stomach wall, **(B)** membrane ionic currents and intracellular Ca^2+^-IP_3_ components included in an interstitial cell of Cajal (ICC) model, and **(C)** membrane ionic currents in a smooth muscle (SM) cell model. See **Table 1** for symbols and details of ionic currents. [Ca]_i_ = Intracellular Ca^2+^, ER= endoplasmic reticulum, Na^+^/Ca^2+^ = Sodium/Calcium exchange pump, SR = Sarcoplasmic reticulum, SS = Submembrane space, **(D)** Simulated rhythmic membrane potential dynamics in an ICC (left panel) and an SM cell (right panel), respectively.

The gastric motility network architecture consists of a longitudinal chain of ICCs bordered by a similar inner chain of SM cells, with one-to-one connectivity formed through nearest-neighbor coupling as shown in **Fig 2**; This closely mimics their biological arrangement in the mammalian stomach [14,39,45]. Endogenously ICCs are coupled via gap junctions [15–17] which permit exchange of ions and other small molecules; We therefore incorporated both an electrical conductance and second messenger permeabilities, namely, Ca^2+^ and IP_3_, between ICCs [22,23]. Between each ICC and its connected SM cell, and between adjacent SM cells we incorporated electrical conductance-based coupling [15]. For the overall network, we incorporated 42 ICCs and 42 SM cells formulated using >1500 ODEs and >2000 parameters. As such the network offers a complex, but realistic, framework to evaluate mechanisms involving ICC-ICC, SM-SM, and ICC-SM coupling and their contributions to inter-ICC coordination resulting in the GSW propagation.

**Table 1:**
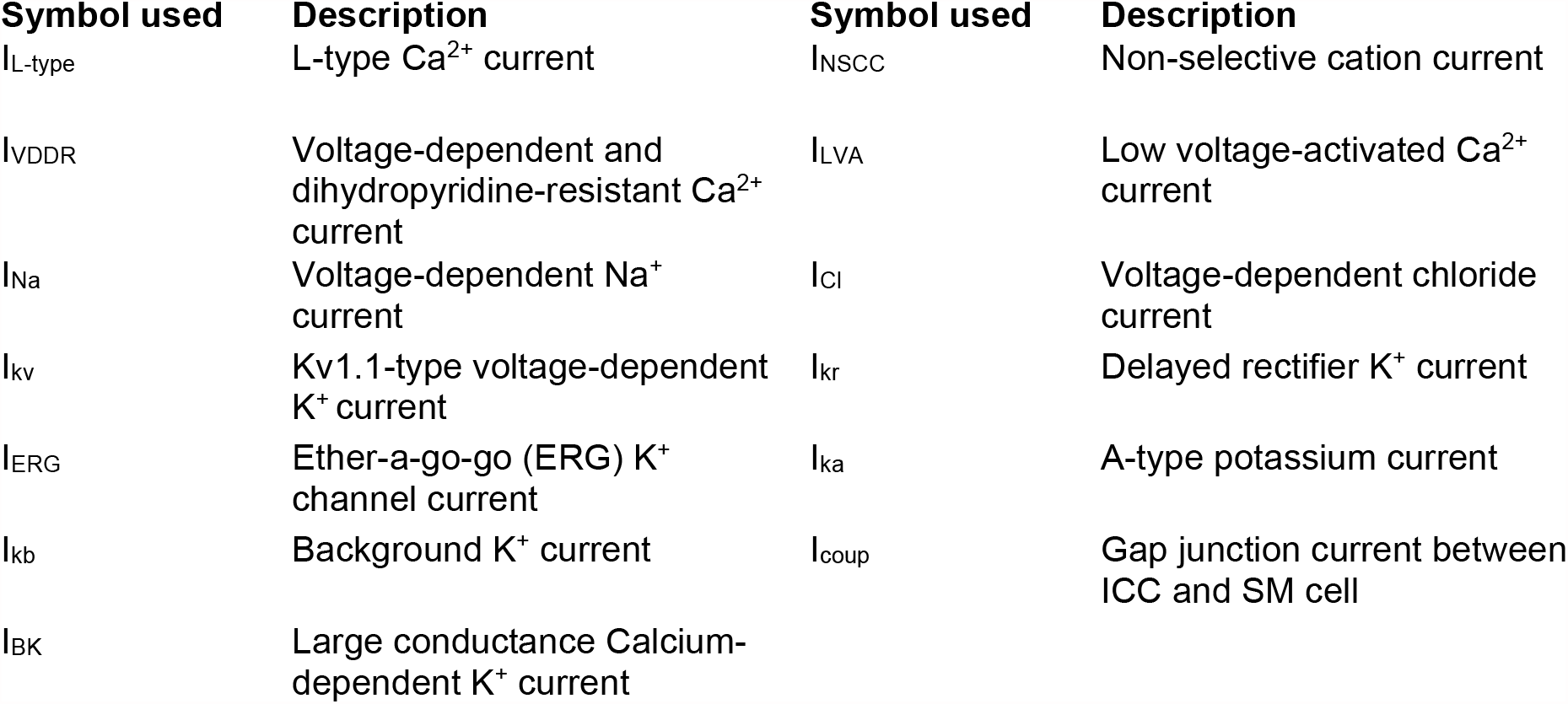
Ionic currents in the ICC model.

**Fig 2.**
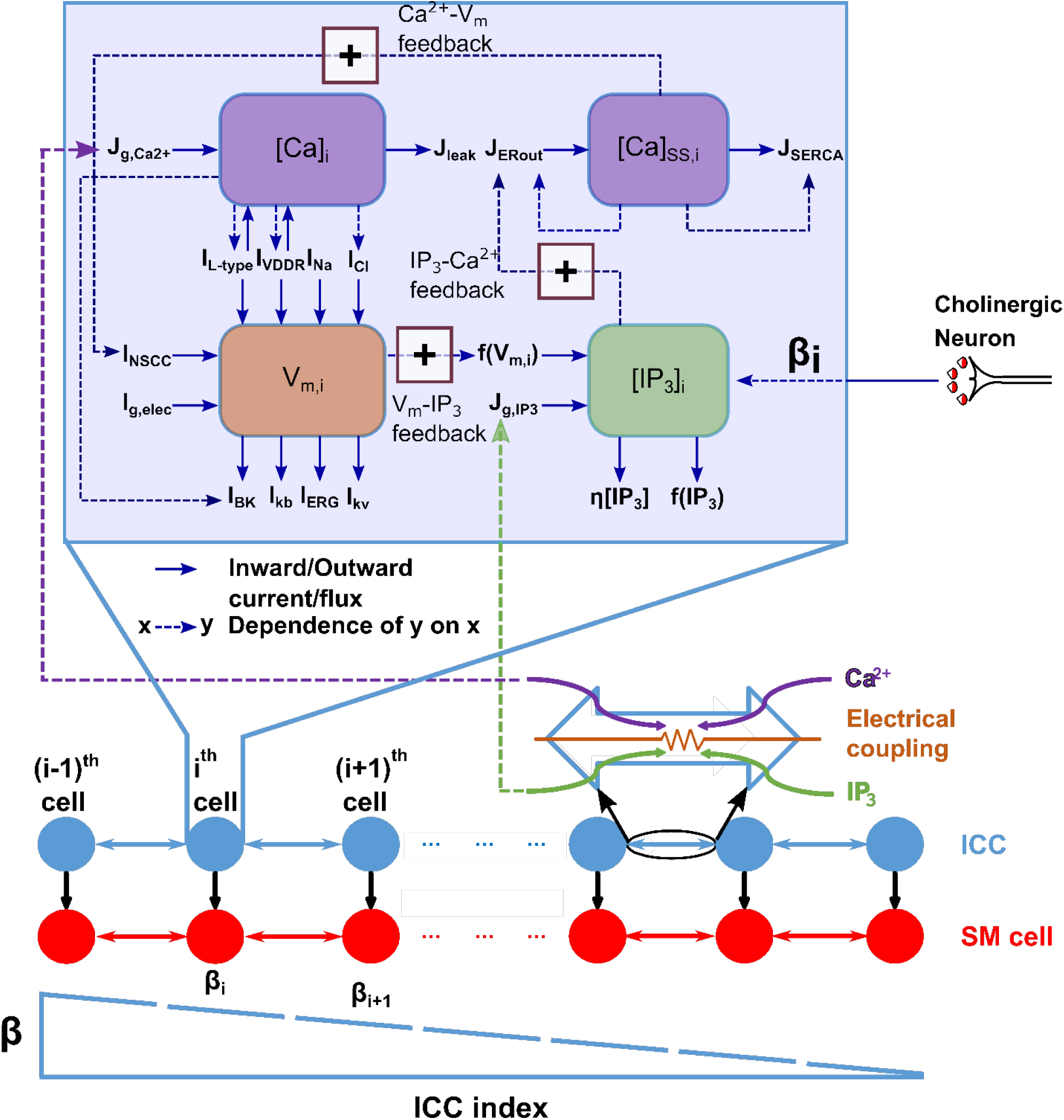
Gastric Motility Network model architecture. The GMN is constituted of a chain of nearest-neighbor coupled interstitial cells of Cajal (ICC) and associated smooth muscle (SM) cells. The ICCs have electrical and second messenger (Ca^2+^ and IP_3_) based coupling, while the ICC to SM and SM to SM couplings are only electrical. There is a negative gradient in enteric neural stimulus (β) that modulates IP_3_ production rate along the rostrocaudal ICC chain (ICC index). Key variables and their interconnections impacting the functionality of an ICC are illustrated in the enlarged inset. The bold arrows in the inset show the inflows and outflows of key variables. The dashed arrows indicate dependency of the sink variable on the source variable. The dashed arrows with (+) sign highlight the positive feedback pathways important for intrinsic pacemaking by an ICC. See **Table 1** and text for further details.

**Fig 2** highlights the framework used to assess whether and how the intrinsic and extrinsic mechanisms engaging second messengers control the inter-ICC coordination necessary for generation of the GSW and thus, the *entrainment range* in this GMN. The enlarged box inset shows the intracellular pathways of the ICC model essential for its intrinsic oscillatory behavior or pacemaker activity. These consist of the multi-stage feedback pathway between the membrane potential, *V*_*m*_ and the intracellular Ca^2+^ and IP_3_ concentrations. The intracellular Ca^2+^ is further divided into two compartments: Ca^2+^ in the sub-space (SS) consisting of the ER and mitochondria ([*Ca*]_*ss,i*_) and near membrane Ca^2+^ ([*Ca*]_*i*_). The bold arrows in the schematic show the inflows and outflows of the key model variables (see **Table 1** in **Methods** for description of the membrane ion channel currents). The dashed arrows with annotation highlight the positive feedback pathways important for intrinsic pacemaking in the ICC model. Note that the membrane potential exerts a positive feedback on intracellular IP_3_ concentration (see *V*_*m*_ − *IP*_3_ feedback in **Fig 2, Equation 4** in **Methods**, also [20,27,41]); The intracellular IP_3_ concentration, [*IP*_3_]_*i*_, in turn affects the [*Ca*]_*ss,i*_ concentration through the outward Ca^2+^ flux from the ER (*J*_*ERout*_) [46–48], as highlighted by the *IP*_3_ − *Ca*^2+^ feedback in **Fig 2**. Lastly, the [*Ca*]_*ss,i*_ increases a non-selective cation channel current (NSCC) which closes the loop by impacting the membrane potential (the *Ca*^2+^ − *V*_*m*_ feedback) [49,50].

An initiating event for pacemaker activity (enteric neural stimulus) is assumed to increase the IP_3_ production rate and is modeled as a constant rate, *β*, for each ICC (see **Equation 4** in **Methods**). Biologically, increases in IP_3_ levels can originate from endogenous release of neurotransmitters from enteric neurons [26] and subsequent activation of G-protein coupled muscarinic receptors on ICCs at cholinergic synapses along the gastrointestinal (GI) tract [26,51]. In our model, we assume the neurotransmitter release and uptake by receptors as a single event termed as ‘enteric neural stimulus’. The enteric neural stimulus driven increase in [IP_3_] results in an increase in the [*Ca*]_*ss,i*_ mimicking the endogenous release of Ca^2+^ from IP_3_ -operated stores in the ER. In response to [*Ca*]_*ss,i*_ increase, the Ca^2+^ uniporter on the mitochondrial membrane is gated open, and Ca^2+^ ions flow into the mitochondria down the steep electrochemical gradient (the term *J*_*Uni*_ in **Equation 3** in **Methods** represents this mechanism). This is thought to remove a larger number of Ca^2+^ ions from the subspace than had previously entered from the ER, causing a temporary drop in the subspace Ca^2+^ concentration [42,49,50]. This activates the non-selective cation channel current leading to membrane depolarization and onset of oscillation in the ICC. Subsequently, further increase in [*Ca*]_*i*_ due to opening of voltage-dependent Ca^2+^ channels is followed by activation of voltage- and Ca^2+-^dependent K^+^ currents to cause repolarization that restores *V*_*m*_ to hyperpolarized values. The above events repeat to cause the regenerative pacemaker potential activity.

To enable longitudinal entrainment of ICCs along the stomach’s length with a rostrocaudal frequency gradient we set a rostrocaudal gradient for the enteric neural stimulus (*α*) that modulates IP_3_ production along the ICC chain (see **Fig 2**). Such a gradient reflects evidence that cholinergic inputs to the ICCs show a rostrocaudal decrement along the GI tract [52,53].

The overall framework allowed us to examine whether engaging adjacent ICCs via two distinct mechanisms would result in similar entrained slow-wave. One mechanism relies on the electrical conductivity of the gap junction coupling between ICCs which *directly* depolarizes their membrane voltage during entrainment. This is a widely used formalism used in previous modeling studies [7,17]. The second mechanism based on exchange of IP_3_ and Ca^2+^ between adjacent ICCs has not been explored in previous models to examine the *entrainment range* (however see [8]). In our GMN model we also assume that exchange of second messengers between ICCs can occur via gap junctions [22–24]; the permeabilities are set for these small molecules (see **Equations 6 and 7** in **Methods**).

### Longitudinal entrainment of ICCs produces a slow-wave for normal peristalsis

We tested whether the GMN model can produce a slow-wave of uniform frequency similar to a rostrocaudally propagating slow-wave for normal peristalsis in the intact stomach (e.g., in the cat [45], in the dog [39], and guinea pig and humans [14]). We also examined whether the SM cells driven by the entrained ICCs exhibit the expected positive and constant time lags between adjacent cells from the rostral to the caudal end of the network [30,33,45,54]. The ICC/SM pair function as a pacemaker unit. We report the membrane potential of SM cells as a proxy for the pacemaker unit activity (also see **S8 Fig**) as is done in experimental work [45,55].

The network was simulated for 900 seconds, where steady-state entrainment was observed approximately after 300 seconds of transient response. **Fig 3** highlights the entrainment and the resulting slow-wave in the GMN at steady-state. In **Fig 3A**, a spatiotemporal map of the membrane potentials of the 42 SM cells demonstrates the rostrocaudal propagation of the slow-wave in the network. The grey shaded regions indicate the up-swing, and the black regions the down-swing in the membrane voltage of the SM cells. A single GSW cycle consists of successive phases of activity from the rostral to the caudal-most pacemaker units. In **Fig 3B**, we highlight the membrane potential of every 7^th^ SM cell in the network. We define and measure *SM Cell Period* as the time between two consecutive peaks in the membrane voltage of an SM cell. An increase or decrease in the *SM Cell Period* reflects bradygastria and tachygastria, respectively. *Relative Lag* is defined as the time difference between the peak membrane voltage of SM_1_ and SM_i_, where *i* = 2,3,4…42, and *Total Lag* as the time between the peak membrane voltages of SM_1_ and SM_42_ and. Slow-wave velocity is inversely proportional to the *Total Lag*, provided that the *Total Lag* value reaches a constant value. At steady-state, *Relative Lags* should reach constant values along the length of the network for a normal slow-wave exhibiting entrainment. Any deviation of *Relative Lag* from constancy reflects decoupling.

**Fig 3.**
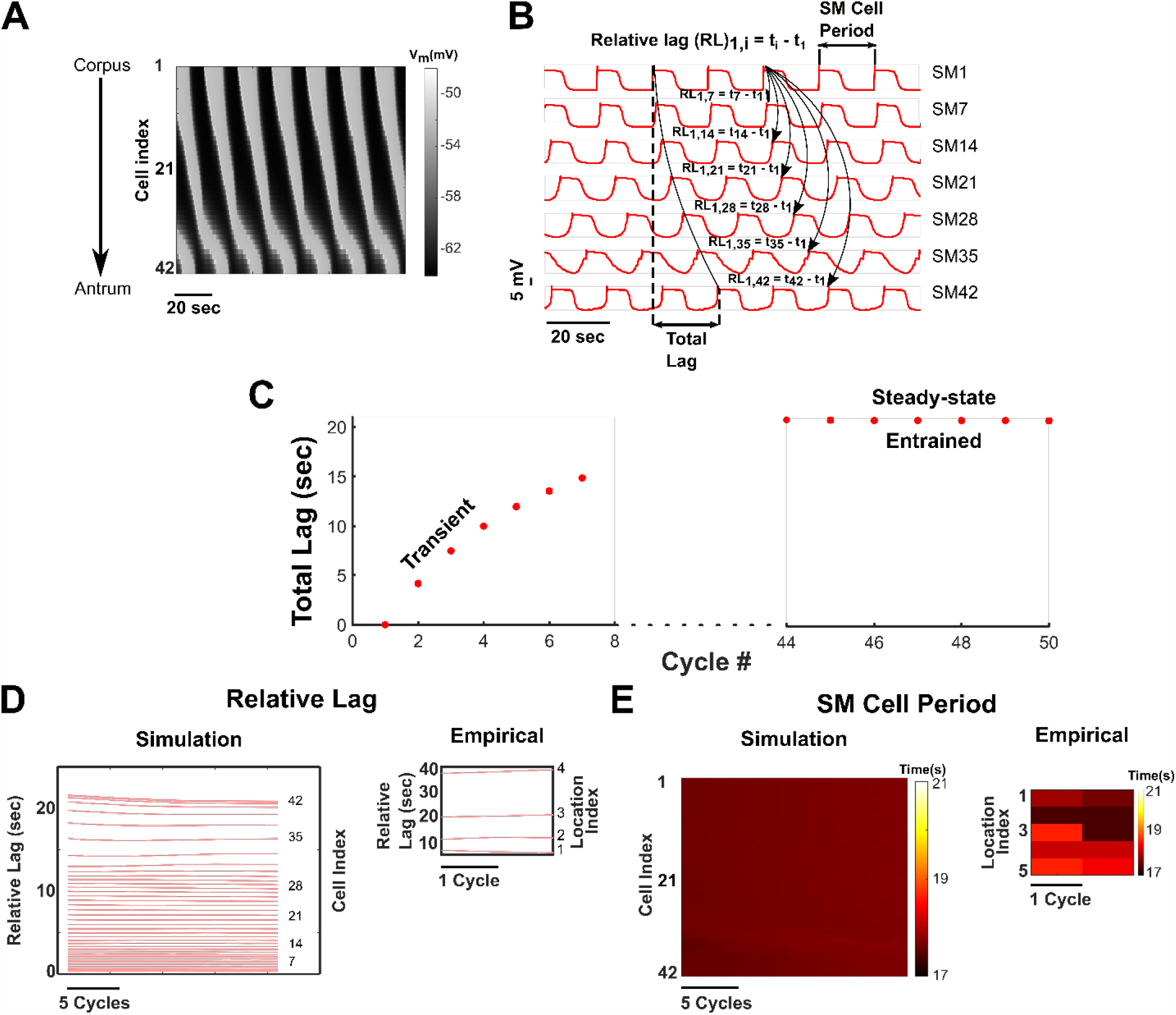
Slow-wave propagation and entrainment in the GMN. **(A)** Spatiotemporal plot of the SM cell membrane potential along the length of the stomach (vertical axis) and time (horizontal axis) of all 42 SM cells of the network. The direction of slow-wave propagation occurs from the rostral end of the network (representing the mid-corpus) to the caudal end (representing the terminal antrum). **(B)** The membrane potential of 7 equidistant SM cells in the 42-cell network. **(C)** *Total Lag* for the first 7 cycles of simulation and the last 7 cycles of simulation (at steady state). **(D)** *Relative Lag* (RL) between the 1^st^ SM cell and i^th^ SM cell in the network for the last 20 cycles, where i = 2, 3, 4,…, 42. The lag increases from the rostral end of the network to the caudal end, indicating a decrease in slow-wave velocity. **(E)** Spatiotemporal map of periods of all 42 SM cells in the network for the last 20 cycles. The inset plots for the *Relative Lag* (**Fig 3D**) and *SM Cell Period* (**Fig 3E**) were generated from the *in vitro* recordings of SM activity at different locations along the length of the cat stomach [45], where location 1 is the rostral-most recording.

In our model, at steady-state, the average *SM Cell Period* over 7 slow-wave cycles for the boundary cells SM Cell 1 and SM Cell 42 were 17.7 (± s.d. = 0.01) seconds and 17.7 (± s.d. = 0.01) seconds respectively. These periods match closely with those observed *in vitro* in studies of cat [45] and guinea pig stomach SM cells [14]). At steady-state, the computed *Total Lag* reached an approximately constant value of 20.8 (± s.d. 0.02) seconds (**Fig 3C**), and it was consistent between consecutive cycles. The *Relative Lag* for each cell reached an approximately constant value (**Fig 3D**) as well as the periods for all the SM cells in the network (observed from the spatiotemporal map of *SM Cell Periods* in **Fig 3E**). These results indicate the entrainment of all the pacemaker units to a uniform gastric slow-wave frequency.

For comparison with empirical results, we also performed meta-analysis of *in vitro* recordings from the cat stomach reported by *Xue S et al*., [45] and generated the *Relative Lag* and spatiotemporal map of *SM Cell Periods*. These are illustrated as insets in **Fig 3D** and **3E** and corroborate our model findings. Thus, we demonstrate that our GMN model is a biologically plausible comprehensive network capable of generating an entrained slow-wave in the stomach with the appropriate rostrocaudal phase lags to support normal peristalsis.

### Inter-ICC electrical coupling and second messenger exchange synergize to generate slow-wave propagation

Our *in silico* model with distinct electrical gap junction conductance and IP_3_ and Ca^2+^ permeabilities enabled us to study their specific contributions to network entrainment and GSW properties as shown in **Fig 4**. We examined the behavior of the *in silico* GMN model with coupling between ICCs through electrical gap junctions alone (**Fig 4A1**), with coupling through second messenger exchange of Ca^2+^ and IP_3_ (**Fig 4A2**) alone, or with both (**Fig 4A3**) over 900 seconds (as for the default network in **Fig 3**). When either electrical gap junction coupling or Ca^2+^ and IP_3_ permeabilities alone were present, we observed partial entrainment along the rostrocaudal chain as demonstrated by the *Relative Lags* (**Fig 4B1** and **4B2**) and the *SM Cell Periods* (**Fig 4C1** and **4C2**) and the results with electrical gap junction coupling alone (**Fig 4B1** and **4C1**) were akin to those observed in motility disorders where slow waves are decoupled and bradygastria is observed in the antrum while the corpus continues to demonstrate normal electrical activity [56]. However, when both constitutive mechanisms were activated simultaneously, an enhanced *entrainment range* (**Fig 4A3, 4B3** and **4C3**) was obtained suggesting that addition of exchange of second messengers to electrical coupling has a synergistic effect.

**Fig 4.**
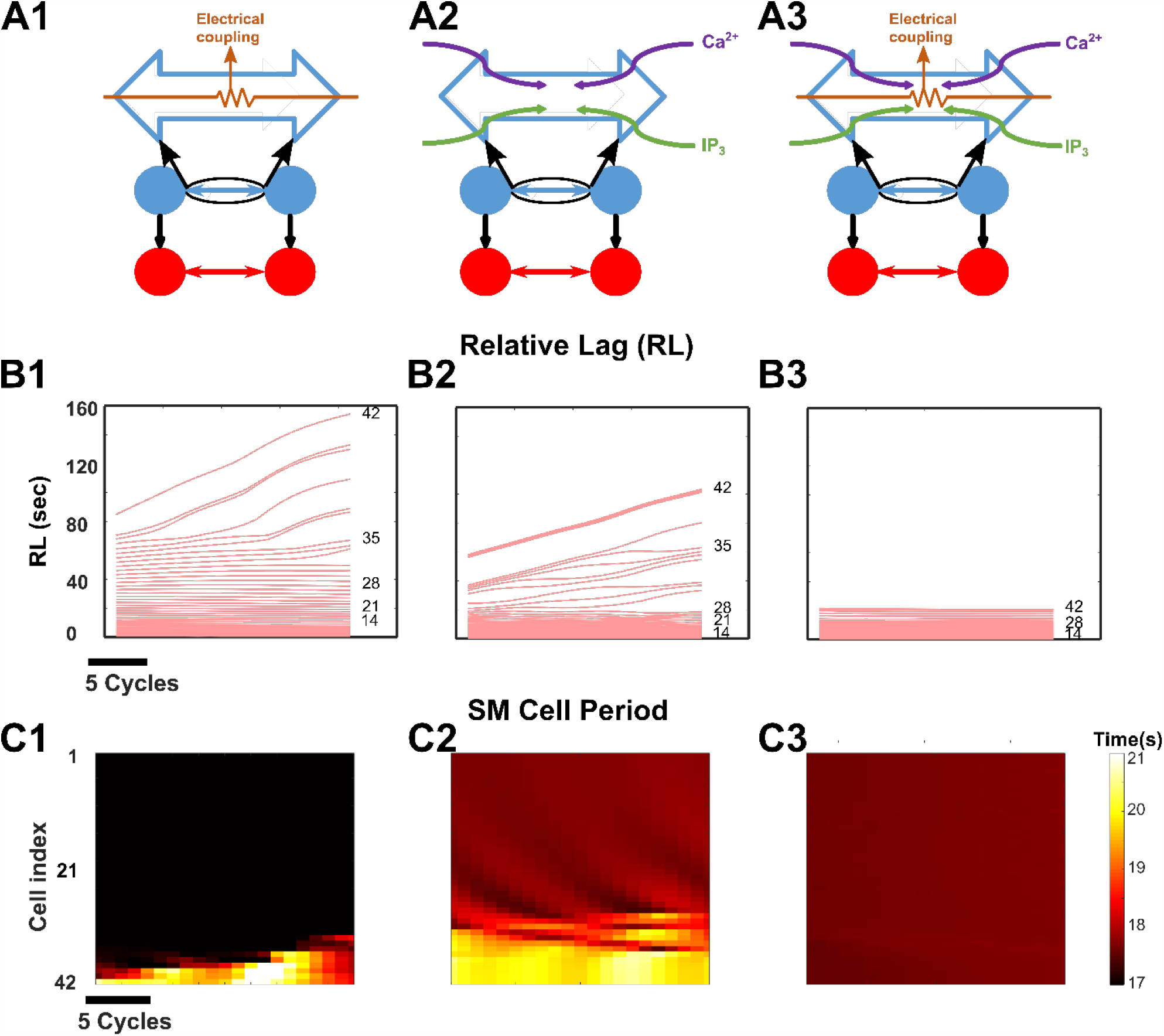
**Enhanced longitudinal entrainment. Synergy of electrical coupling and second messenger exchange preserve longitudinal entrainment along the entire length of the stomach. (A1, A2 and A3)** Structure of the network, when only electrical coupling is present **(A1)**, when only exchange of second messengers is present **(A2)**, and when both electrical coupling and exchange of second messengers are present **(A3). (B)** The *Relative Lags* for the three different cases shown in **Fig 4A** are measured from the last 20 cycles of respective simulations. **(C)** Spatiotemporal maps of *SM Cell Periods* in the three different cases respectively.

### Impact of increasing strengths of electrical gap junction coupling and exchange of second messengers between ICCs

We performed a sensitivity analysis to characterize the effects of increasing strengths of electrical conductivity versus second-messenger permeabilities in inter-ICC coupling on network entrainment and found that they influence the entrainment range, the gastric slow-wave velocity, and the pacemaking frequency. **Fig 5A-5C illustrate** the network’s behavior for increasing inter-ICC electrical coupling conductance, *G*_*ICC*− *ICC*_ two times above and below a default value of 0.7 nS. The spatiotemporal maps of SM cell membrane potential and the corresponding *Relative Lag* and action potential periods of SM cells are respectively shown for the lowest (**Fig 5B1-5B3, left**) and highest (**Fig 5B1-5B3, right**) *G*_*ICC*− *ICC*_ values considered. In this range of coupling conductance strengths, the network transitioned from partial entrainment (**Fig 5B1-5B3, left**) to complete rostrocaudal entrainment (**Fig 5B1-5B3, right**). For the lowest *G*_*ICC*−*ICC*_ value considered, the *Relative Lag* diagram illustrated in **Fig 5B2, left** shows decoupled slow waves, whereas the *SM Cell Period* diagram depicted in **Fig 5B3, left** demonstrates signs of bradygastria. Steady-state rostrocaudal entrainment was achieved only for higher values of the *G*_*ICC*−*ICC*_ as shown by the levelling out of *Total Lag* in the color-coded insets in **Fig 5C**. Overall, an increase in the strength of electrical coupling conductance produced an enhancement in the *entrainment range*.

**Fig 5.**
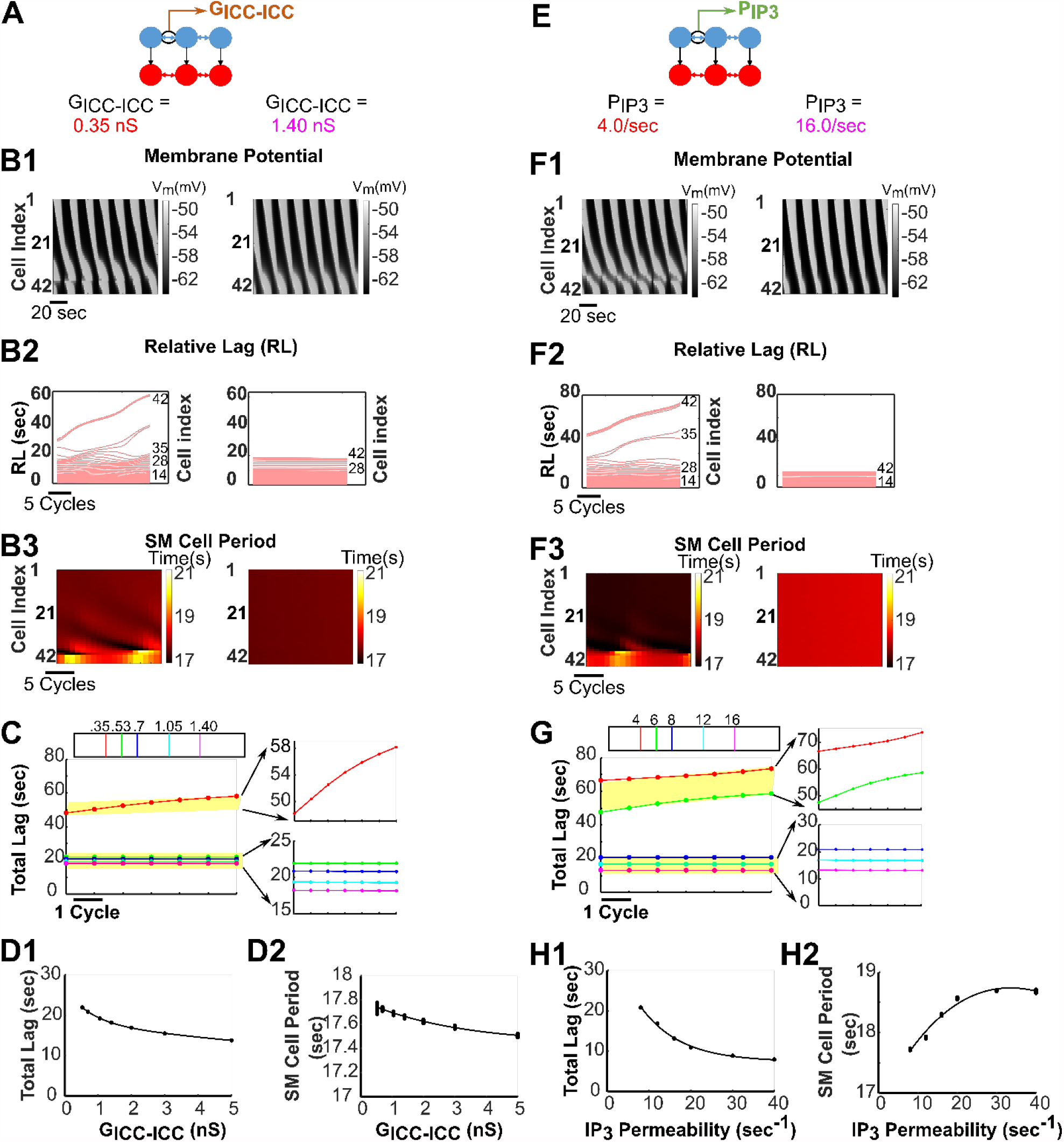
**Electrical gap junction and second messenger coupling strengths. The inter-ICC coupling strengths impact GMN entrainment, gastric slow-wave velocity and pacemaker frequency.** **(A, E)** *G*_*ICC*−*ICC*_ and *P*_*Ip*3_ are altered for the simulations in **Fig 5B-5D** and **Fig 5F-5H**, respectively. **(B)** Spatiotemporal map of membrane potential **(B1)**, *Relative Lags* **(B2)**, and spatiotemporal map of *SM Cell Periods* **(B3**) for the network when *G*_*ICC*−*ICC*_ = 0.35 nS (left panels, partially entrained) and 1.4 nS (right panels, entrained). **(C)** The *Total Lag* for changes in *G*_*ICC*−*ICC*_ is shown for the last 7 cycles of 900-sec simulations for different conductance values (nS) indicated by the color legend. The two distinct classes of responses are enlarged in the right panels (lower ones entrained, upper ones partially entrained). An increase in *Total Lag* indicates decrease in slow-wave velocity. **(F)** Spatiotemporal maps of membrane potential **(F1)**, *Relative Lags* **(F2)**, and spatiotemporal maps of *SM Cell Periods* **(F3)** for the network when *P*_*IP*3_ = 4.0 sec^-1^ (left panels, partially entrained) and 16.0 sec^-1^ (right panels, entrained). **(G)** The *Total Lag* for changes in *P*_*IP*3_ is shown for the last 7 cycles of 900-sec simulations for different conductance values (nS) indicated by the color legend. **(D, H)** For several networks, the mean *Total Lag* (odd numbered panels) and the *SM Cell Period* (even numbered panels) of the last 7 cycles for each network with respect to its *G*_*ICC*−*ICC*_ and *P*_*IP*3_ can be fit by individual exponential function, respectively. For increasing values of *G*_*ICC*−*ICC*_, an exponential fit (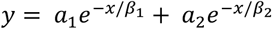, where *C*_1_ = 8.65, *α* = 0.75, *C*_2_ = 18.23, *α*_2_ = 17.87) has been drawn along the mean value of *Total Lag* and for increasing values of *P*_*IP*3_, another exponential fit (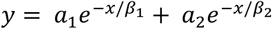, where *C*_1_ = 7.44, *α* = 1.70e5, *C*_2_ = 32.07, *α*_2_ = 9.37) has been drawn along the mean values of *Total Lag*. Partially entrained networks have not been considered for equation fitting. For *SM Cell Period* calculation, the last cell (42^nd^ cell) has been considered as the representative cell of the network. An increase in *Total Lag* indicates decrease in slow-wave velocity while a decrease in *SM Cell Period* indicates increase in pacemaking frequency.

**Fig 5E-5G** present the network outcome for increasing IP_3_ permeability (values of *P*_*IP*3_ two times above and below a default value of 8.0 sec^-1^) at the default value for the electrical coupling conductance. The spatiotemporal maps of SM cell membrane potential and the corresponding *Relative Lag* and action potential periods of SM cells are shown for the lowest (**Fig 5F1-5F3, left**) and highest (**Fig 5F1-5F3, right**) *P*_*IP*3_ values considered. Similar to increasing *G*_*ICC*−*ICC*_ values, the networks with increasing *P*_*IP*3_ values demonstrated a transition from partial entrainment to complete rostrocaudal entrainment. For the lowest *P*_*IP*3_ value considered, the network generates decoupled slow waves as shown in the *Relative Lag* diagram of **Fig 5F2, left**; however, unlike lowest *G*_*ICC*−*ICC*_ value, the network having lowest *P*_*IP*3_ value considered demonstrates signs of tachygastria as illustrated by the *SM Cell Period* diagram in **Fig 5F3, left**. The tachygastria can be prevented by increasing exchange of IP_3_ across ICCs (**Fig 5F3, right**) [57].

For increasing *G*_*ICC*−*ICC*_ and *P*_*IP*3_, in both cases, the effect on *Total Lag* was qualitatively similar, i.e., decreased with increasing coupling strengths and reached an asymptote as shown in **Fig 5D1 and 5H1** (actual values are provided in **S2, S4 Tables** respectively). These results suggest that increasing inter-ICC coupling strengths leads to robust rostrocaudal network entrainment wherein the gastric slow-wave velocity approaches a limit.

However, *G*_*ICC*−*ICC*_ and *P*_*IP*3_ increases differentially altered the *SM Cell Periods* in the entrained networks at steady-state. This is shown in **Fig 5D2** and **Fig 5H2** respectively (actual values in **S3** and **S5 Tables**). The *SM Cell Period* decreased with increasing *G*_*ICC*−*ICC*_, whereas increasing values of *P*_*IP*3_ increased the *SM Cell Period*. These results suggest that with an increase in *P*_*IP*3_, a decrease in the pacemaking frequency occurs whereas the gastric slow-wave velocity increases. Collectively, the results suggest that on an increase of electrical coupling strength and IP_3_ permeability, the velocity of gastric slow-wave propagation increased while the pacemaker potential frequency increased with the former and decreased with the latter.

We also exclusively increased the Ca^2+^ permeability (*IP*_*C*2+_) to test its effects on network entrainment. We noted that even two orders of magnitude changes above and below the default value of *IP*_*C*2+_ produced negligible effect on all the network characteristics analyzed: *entrainment range, Relative Lag, SM Cell Periods* and *Total Lag* in the entrained networks (see **S6 Fig**). This is likely due to an order of magnitude lower permeability of Ca^2+^ compared to IP_3_ for gap junctions [18]. Due to this lack of effect of *IP*_*C*2+_ on entrainment and gastric slow-wave characteristics, in what follows, we will only focus on changing *P*_*IP*3_.

### Inter-ICC IP_3_ exchange preserves longitudinal entrainment

Variability in ICC coupling strengths could stem from variable gap junction densities in the stomach [58,59]. We find that inter-ICC IP_3_ ensures appropriate slow-wave propagation and stable pacemaking potential generation under variable ICC coupling strengths. We first introduced variability in coupling conductance by random sampling of *G*_*ICC*−*ICC*_ values from a uniform distribution (see corresponding results in **Fig 6A-6F**) in the absence of inter-ICC IP_3_ exchange. The range of conductance values were *default* ± 20% or ± 50% or ± 100% (see **Fig 6A**). As shown in **Fig 6B**, the *Total Lags* have not reached a steady-state value in any of the three networks. Consequently, slow-wave velocity cannot be inferred from here and the failure of *Total Lag* to attain a constant value indicates absence of rostrocaudal network entrainment. **Fig 6C** shows the distribution of *Total Lags* computed over the last 7 cycles of simulation, using violin plots. From these plots we note that the variability in *Total Lags* consistently increases with the variability in coupling conductance strength. The SM cell membrane potential (**Fig 6D**), *Relative Lag*, (**Fig 6E**) and *Period* (**Fig 6F**) indicate that variability of *G*_*ICC*−*ICC*_ indeed gave rise to clusters of partial entrainment. The *Relative Lag* diagrams in **Fig 6E**, in particular, provide evidence for the decoupled slow waves. The *SM Cell Period* diagrams in **Fig 6F1 and 6F2** bears close resemblance to bradygastria whereas **Fig 6F3** demonstrates a mix of bradygastria and tachygastria. These results suggest that, in the absence of inter-ICC IP_3_ exchange, decoupled slow waves and bradygastria and/or tachygstria can coexist.

**Fig 6.**
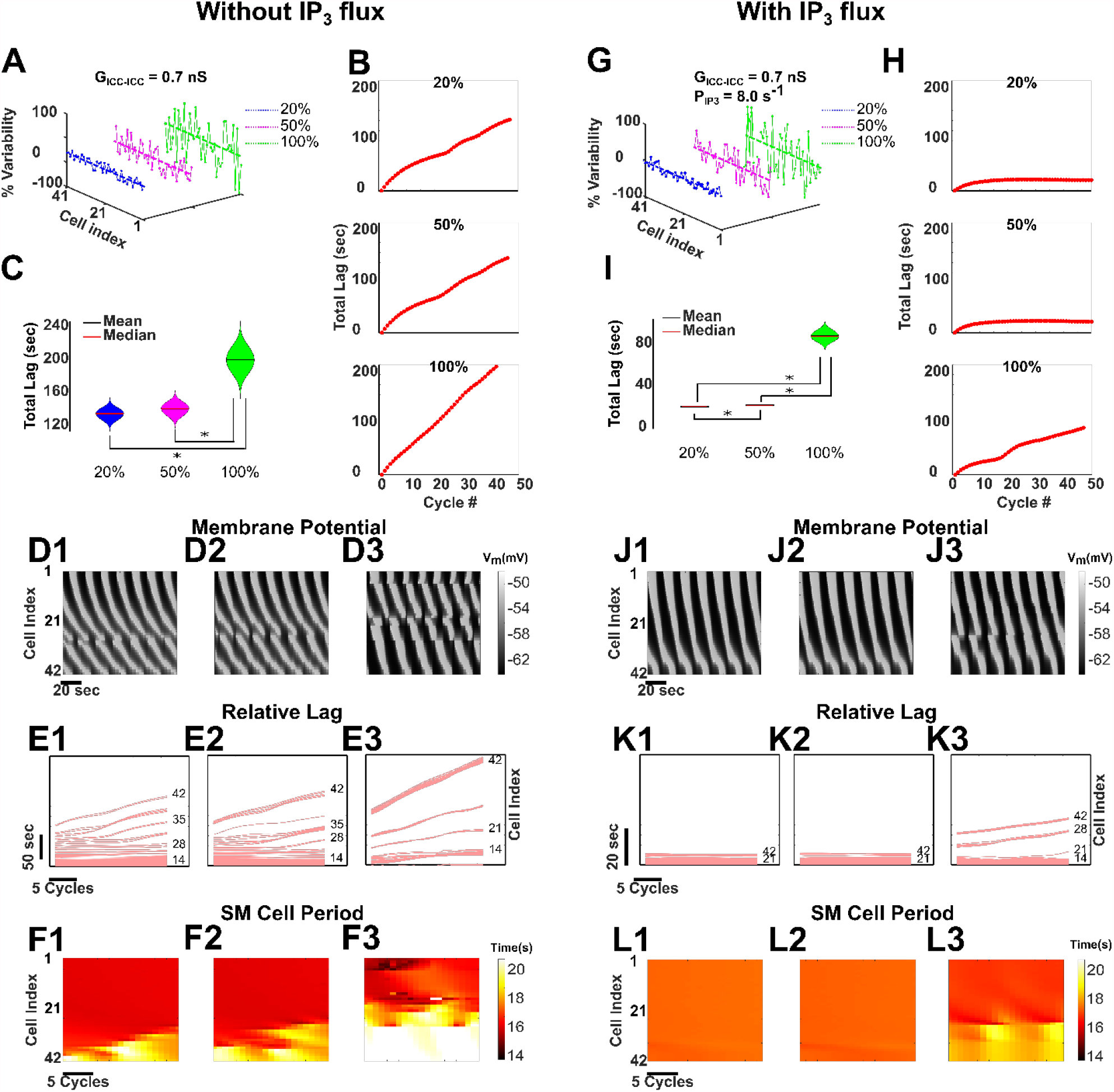
Entrainment in GMN is resilient to variability in inter-ICC coupling strengths. **(A**,**G)** The variability in only *G*_*ICC*−*ICC*_ (**A**) and both *G*_*ICC*−*ICC*_ and *P*_*IP*3_ (**G**) (20%, 50%, and 100% variability) are shown around their mean values. **(B, H)** *Total Lag* of the network without inter-ICC IP_3_ exchange for three different variabilities in *G*_*ICC*−*ICC*_ **(B)** and of the network with inter-ICC IP_3_ exchange for three different variabilities in *G*_*ICC*−*ICC*_ and *P*_*IP*3_ **(H)**. The latency to entrainment cannot be inferred from any of the panels in **(B)**, since the *Total Lag* has not reached steady-state value in any of them. However, the latency to entrainment can be measured from the first two panels in **(H)**, since the *Total Lag* has attained constant value in these two cases. **(C, I)** *Total Lag* is shown for each case for the last 7 cycles in steady-state. A violin plot shows the differences in *Total Lag* for all three cases. The asterisk (*) symbol represents a statistically significant difference between the corresponding quantities. Spatiotemporal diagrams of membrane potential **(D, J)**, *Relative Lags*, **(E, K)** and spatiotemporal maps of *SM Cell Periods* **(F, L)** for the network are shown for three different variabilities in *G*_*ICC*−*ICC*_ for the network without inter-ICC IP_3_ exchange and in *G*_*ICC*−*ICC*_ and *P*_*IP*3_ for the network without inter-ICC IP_3_ exchange, respectively.

Next, we included the inter-ICC IP_3_ exchange along with electrical coupling and varied both the *G*_*ICC*−*ICC*_ and *ICC*_3_ values 20%, 50%, and 100% about their corresponding default values of 0.7 nS and 8.0 sec^-1^, respectively (**Fig 6G**). Interestingly, addition of IP_3_ exchange enabled rostrocaudal entrainment for networks with 20% and 50% variability in these coupling parameters, but not with 100% variability (see panels in **Fig 6H**). The distribution of *Total Lags* in **Fig 6I**, the SM cell membrane potential (**Fig 6J**), *Relative Lag* (**Fig 6K**), and *Period* (**Fig 6L**) further confirms these results. The *Relative Lag* diagrams in **Fig 6K1 and 6K2** show absence of decoupling for 20% and 50% variability, respectively; in contrast, for 100% variability, **Fig 6K3** shows decoupled slow waves as evidenced by temporal changes in *Relative Lag*. Parallel observations have been made from the *SM Cell Period* diagrams in **Fig 6L1 and 6L2**, where signs of abnormal rhythmicity (bradyagastria or tachygstria) are effectively eliminated, although for 100% variability, there are still signs of bradygastria (**Fig 6L3**). Furthermore, **Fig 6H** demonstrates that there is no substantial difference in the latency to reach entrainment between the networks having 20% and 50% variability. Together these results suggest that the existence of intercellular exchange of IP_3_ can preserve the longitudinal entrainment to a greater extent along the length of the stomach, thereby preventing decoupling and restoring the normal behavior.

## Discussion

We have developed a computational framework consisting of a novel non-linear mathematical model for slow-wave propagation in the stomach wall that includes physiologically established intra- and intercellular mechanisms. Utilizing this framework, we have assessed the contribution of intercellular electrical coupling and intercellular exchange of second messengers on longitudinal entrainment of the gastric slow-wave. The intracellular concentrations of Ca^2+^ and IP_3_ are modulated by intercellular exchange of respective molecules (see **Equations 2 and 4**). We found that our model with dynamically coupled nonlinear oscillators fed with second messengers from intercellular exchange and enteric neural stimuli, can regulate the frequency of contractions and velocity of the slow-wave, and can enhance the range of longitudinal entrainment of the gastric slow-wave. The combination of electrical coupling and exchange of second messengers provides robustness to the *entrainment range* in the presence of biological variability. In summary, our detailed analyses of the ICC coupling mechanisms reveal ways in which the constitutive gap junction coupling mechanisms can enhance the network range and stability of the gastric slow-wave essential for peristaltic movement of food and fluid. Furthermore, this model can be used to examine novel hypotheses concerning aberrant mechanisms that may underlie different motility disorders.

### Development of Gastric Motility Network (GMN) model

In the present study, we developed a computational model consisting of a longitudinal arrangement of biophysically based ICC and SM cells found in the stomach wall. Since ICCs and their network are regarded as the key players for generation of pacemaker potentials and propagation of slow waves, we investigated whether and how the intrinsic properties of ICCs that are also modulated by enteric neural inputs enable inter-ICC coordination essential for entrainment. We also examined the crucial intercellular pathways that influence the *entrainment range*, important for long distance GSW propagation necessary for normal peristalsis. For this, we considered multiple interacting feedback pathways involving the intracellular key variables such as membrane potential, Ca^2+^ and IP_3_ concentration of an ICC as well as intercellular electrical coupling and exchange of second messengers. Further based on corroborative evidence that enteric neuron inputs to ICCs show a rostrocaudal gradient [52,53], and indirect evidence that indicates that neurotransmitters enhance IP_3_ in a variety of pacemaker cells [27– 29], we assumed a gradient in the enteric neural stimulus along the ICC chain. Such a gradient was essential for the rostrocaudal entrainment of the slow-wave as also demonstrated in previous computational models [7,19].

### Control of slow-wave characteristics by inter-ICC IP_3_ exchange

First, our simulation results for single ICC and SM cells agree qualitatively and semi-quantitatively with the experimentally measured results (**Fig 1**), thus validating the GMN model components. Next, our results show that the GMN with its multi-stage feedback between IP_3_ -Ca^2+^-V_m_ combined with the intercellular pathways (**Fig 2**) can generate slow-wave propagation with a uniform frequency and a uniform rostrocaudal lag, i.e., entrainment (**Fig 3**). Particularly, the presence of inter-ICC IP_3_ exchange in addition to the gap junction electrical conductance can increase the range of longitudinal entrainment (**Fig 4**), while simultaneously minimizing the signs of bradygastric decoupled slow waves. Our observations are in accordance with the suggestion that bradygastria can potentially result from a failure of normal entrainment [60]. While the distinct intercellular mechanisms modeled in our network (exchange of second messengers and electrical coupling) are experimentally inseparable, the biophysical formulation included in the GMN model allowed examination of their individual contributions to longitudinal entrainment of ICCs. Second, we systematically evaluated the network behavior for increasing values of electrical conductance and IP_3_ permeability (since Ca^2+^ permeability was shown to have little to no impact on GSW entrainment). Surprisingly, the IP_3_ exchange affected the features of the slow-wave in a manner that can stabilize the propagating wave (from tachygastric decoupled slow waves for low IP_3_ to a normal coupled gastric slow-wave for elevated IP_3_). In particular, increased IP_3_ permeability of gap junctions resulted in an *increase* in the *SM Cell Period* (bordering the signs of bradygastria) whereas increased gap junction electrical conductance caused a *decrease* in the *SM Cell Period* (bordering the signs of tachygastria). Thus, in combination with the contrasting effects of electrical conductivity, the IP_3_ coupling could flexibly modulate the SM cell frequencies. We suggest that the apparent compensatory effect of IP_3_ permeability could act as a brake on preventing runaway tachygastria (elevated SM cell frequencies, which is inversely related to *SM Cell Period* or pacemaking potential duration) in the event of increasing electrical coupling strengths. Increased gap junction density, which is reflected by increased gap junction coupling strength in our model, has been demonstrated in colonocytes during bacterial infections resulting in diarrhea generation [61]. Whether the same holds true for gastric motility disorders requires further experimental investigation. Even if the electrical gap junction density becomes exceedingly high, the inter-ICC IP_3_ coupling makes sure that the SM cell frequency, and therefore, gastric contraction frequency stays in a physiologically plausible range.

Interestingly, increasing values of electrical conductance or IP_3_ permeability led to an increased slow-wave velocity. In our results, the increase of slow-wave velocity was demonstrated by a decrease in the *Total Lag* of the slow-wave (**Fig 5**). These observations are very similar to the observations in studies of the dog small intestine where a negative gradient of gap junctions exists across the duodenum, jejunum and ileum and so does the slow-wave velocity in these compartments of the small intestine [62]. Electrophysiological experiments with gap junction enhancers (such as in Rotigaptide [63]) also support the observations of our computational study. As per our knowledge, such gap junction enhancers have not been employed as medicinal interventions yet, although they have the potential to elevate the reduced slow-wave velocity as observed in gastroparesis patients [60]. We would have to be very cautious though in introducing such enhancers, since slow-wave velocity has been reported to be already highly elevated in the gastric corpus of aged patients having gastroparesis with impaired peristalsis [64] and somewhat elevated in glucagon induced hyperglycemic dogs [6]. Future experiments could investigate the proper dosage of the gap junction enhancers so that the slow-wave velocity remains in a moderate range. For now, we can only speculate that in an intact stomach, the electrical gap junction coupling strength probably does not reach an exceedingly high value, and even if it has a moderately high value, **Fig 5D1 and 5H1** show that the slow-wave velocity reaches a limit, which is within a physiologically reasonable range.

### Distinct period-velocity relationships exist for increasing *G*_*ICC*−*ICC*_ and *P*_*IP*3_

According to coupled oscillator theory, the direction of the slow-wave would depend on the frequency gradient of component pacemaker ICCs, whereas the slow-wave velocity would depend on the intercellular coupling strength [65]. Therefore, ideally, there should be a dispersion relationship between slow-wave velocity and frequency (expressed by *SM Cell Period*, in our case). However, previous studies of the canine gastric antrum [65] and porcine gastric corpus and antrum [66] suggest a dependence of slow-wave velocity on the observed *SM Cell Period*. The period-velocity relationship as noted in **Fig 5H1 and 5H2** (higher the period, higher the velocity) for increasing permeability of IP_3_ supports these *in vitro* [65] and *in vivo* [66] findings. In contrast, the period-velocity relationship noted in **Fig 5D1 and 5D2** (lower the period, higher the velocity) for increasing electrical coupling strength is somewhat surprising. Because, if the period decreases, the velocity of a slow-wave should drop as a response to encroachment on the tail of the previous slow-wave (see **Fig 7 in [66]** for a more intuitive understanding). Although such a relationship has been observed along the length of the intestine [62] under normal conditions, its occurrence is probably due to the emergence of frequency plateaus, which in fact reflect localized decoupling due to the reduced gap junction density at certain points along the intestinal length [59]. Frequency plateaus are not observed in the stomach under normal conditions. Consequently, localized decoupling due to the reduced gap junction density along the length of the stomach is an unlikely explanation for the observed period-velocity relations in **Fig 5D1 and 5D2**. We believe that the less pronounced effect of increasing *G*_*ICC*−*ICC*_ on velocity (reflected as *Total Lag* in **Fig 5D1**) and period (illustrated in **Fig 5D2)** compared to the effect of increasing *P*_*IP*3_ on the same quantities (**Fig 5H1 and 5H2**) makes sure that the trajectory of a slow-wave having increased *G*_*ICC*−*ICC*_ does not appear synchronously with the tail of the previous slow-wave, thus avoiding the positive correlation between period and velocity. In our model results, we are concerned only with the velocity from the rostral end to the caudal end. Measurement of velocity gradient (if there is any) and a non-uniform modeling of gap junction conductance [15] along with the measurement of gap junction density in various animal models could shed further light on the model predictions.

### Exchange of second messengers ensures robust rostrocaudal entrainment when biological variability is present

Finally, we showed that while variability in electrical coupling results in loss of rostrocaudal entrainment (signs of bradygastria and mix of bradygastria and tachygastria observed), presence of IP_3_ exchange despite variability in its permeability, can restore rostrocaudal entrainment (**Fig 6**) and therefore, is essential for peristalsis. Here, inter-ICC IP_3_ exchange offers resilience to the increased variability in coupling strength. This is noteworthy because gap junctions in ICC networks can have variable density in different compartments of the stomach [15,16] and the observed consequences of increasing the variability in inter-ICC coupling in our simulations highlight that this might be a feature important for regulating the wave velocity while maintaining the range of entrainment. Our model with the distinct intercellular mechanisms (exchange of second messengers and electrical coupling) in combination with the intracellular feedback pathways and rostrocaudal neural stimulus gradient allows SM cells to oscillate with a moderate range of frequencies and the gastric slow-wave to propagate with a broad range of velocities (**Fig 5**). Thus, the GMN confers high robustness to the rostrocaudal entrainment of ICCs, including under instances of biological variability in coupling pathways (**Fig 6**). Robustness is a fundamental property of evolvable complex biological systems [67]; a simple mechanism cannot handle extreme changes in physical quantities. Our model offers a new integrative framework for conceptualizing GSW propagation and regulation as a robust system of dynamically coupled oscillators fed with second messengers through intercellular exchange.

### Biological and theoretical assumptions of the GMN model

Although numerous biological mechanisms are involved in the orchestration of gastric motility [68], it is well-established that the peristaltic movement of food/liquid is mediated by a propagating wave of smooth muscle contractions along the stomach wall [5,13,14,69]. This so-called gastric slow-wave involves electrical activity transmitted aborally within the SM cells [55,70]. However, the SM cells on their own cannot produce such a regenerative wave [70,71]. The regenerative electrical activity responsible for the GSW is known to originate largely in the intrinsic pacemaker ICCs [5,42]. Different sets of ICCs innervate circular and longitudinal muscle cells (e.g., myenteric ICCs, intramuscular ICCs [5,13]) and all ICCs may not contribute similarly to pacemaker potential generation and GSW propagation [68]. Our ICC cell model is derived primarily from experimental work on myenteric ICCs, which are widely accepted as intrinsic pacemakers involved in the generation of the pacemaker potential required for GSW propagation [5]. We incorporated known membrane properties in the ICCs and SM cell models using the conductance-based formalism and hand-tuned the conductance parameters to match experimental voltage recordings in these cells (**Fig 1C and 1D**) [45]. Although our model can reproduce rather accurately the established properties of a gastric slow-wave, it should be noted that the description of the underlying mechanisms is by no means exhaustive. For example, we have chosen to use a Hodgkin-Huxley formalism for the ion currents whereas Markovian models would allow for a more detailed description of the complex kinetics of processes (activation, deactivation, inactivation, and recovery from inactivation) that the channels exhibit. However, it can be very challenging to meet the information requirements for defining the transitional rate constants of a Markovian model, in addition to higher computational processing demands [72].

Although ICCs are intrinsic pacemakers, the GSW can significantly be impacted by the coupling between them [8,73,74]. Gap junctions are well-established pathways for such coupling [75]. The gap junctions carry various ions and molecules. Some of these (e.g., K^+^) are passively conducted due to the voltage gradients between adjacent cells (the electrical component), while others spread when there are periodic depolarizations in ICCs that generate these molecules in abundance. The latter are typically second messengers such as IP_3_ and Ca^2+^, which involve chemical coupling via connexin channels [22,23,25,35,36]. The diameter of gap junctional pores allows a wide enough path for Ca^2+^ and IP_3_ to move across the cells through gap junction connexins [76]. To model electrical coupling, a simple ohmic conductance has been widely assumed by previous modeling studies [7,8,19,69,77,78], supported by findings that pacemaker current generated in ICCs is transmitted to the SM cells by gap junction channels located between the ICCs and SM cells [15,17]. We additionally assumed a distinct exchange of second messenger (Ca^2+^ and IP_3_) between adjacent ICCs. Second messenger exchanges are modeled in such a way that they enable Ca^2+^ induced Ca^2+^ release and IP_3_ induced IP_3_ release. Although Ca^2+^ induced Ca^2+^ release is a widely assumed phenomenon within a biological cell, IP_3_ induced IP_3_ release has also been suggested as a plausible event [79]. Ca^2+^ permeability was set an order of magnitude lower than that of IP_3_ permeability. This assumption was based on the fact that the range of diffusion of IP_3_ is orders of magnitude higher compared to free Ca^2+^ (24 µm compared to 0.1 µm). IP_3_ also degrades much more slowly than free Ca^2+^ (1 sec vs. 0.00003 sec) [80]. Previous modeling studies of intercellular Ca^2+^ waves in astrocytes [35] and smooth muscle cells [24] support this approach of modeling Ca^2+^ and IP_3_ permeability. The exchange of IP_3_ was modeled as proportional to the IP_3_ concentration gradient between adjacent cells (see **Methods**, also [81]). Although IP_3_ can diffuse within the cell cytoplasm and not necessarily through the gap junctions, we assumed that IP_3_ only moves to the adjacent cells based on the concentration gradient. For **Fig 3-5**, we considered constant values of gap junction coupling parameters and for **Fig 6**, we assumed uniform distribution of gap junction coupling strength, which resembles the gap junction density. Because of the scarcity of data regarding gap junction distribution, we limited our simulations to only uniform distribution.

In a chain of intrinsic pacemakers, oscillators have their own distinct intrinsic frequency. In an arbitrarily long chain of oscillators, entrainment can emerge provided there exists a linear frequency gradient with fixed frequency difference between the ends [82]. Interestingly in the stomach (also throughout the GI tract), it is known that there is a rostrocaudal frequency gradient in the ICC pacemaker cells. Although such gradients may be achieved by numerous intrinsic and network mechanisms [7,19], we assume that such a gradient in intrinsic frequencies can be achieved due to a maintained gradient in [IP_3_] production rate. Presently it is unclear what sets this gradient tone in IP_3_ production rate. One possibility could be IP_3_ produced by activation of muscarinic receptors due to graded distribution of cholinergic neuron inputs in different functional compartments of the GI tract [52,53]. Although peristalsis can occur even without the help of neural excitation [68], under most circumstances, the enteric nervous system, provides excitation of the musculature required for the stomach wall contraction [26,83], primarily via ICCs. Enteric neurons located in the Myenteric Plexus preferentially directly influence ICCs through neurotransmitters released at synapses which then connect to muscle cells via gap junctions. Since ICCs are closer to the nerve terminal endings and have the muscarinic receptors (M2 and M3) that are responsive to the neurotransmitters, the effect of the Myenteric neurons on ICCs is more important (much smaller gap for neurotransmitters to diffuse) than the direct link to the SM cells (far away and with lesser innervation). Acetylcholine, the primary neurotransmitter released from enteric neurons, is broken down by Acetylcholine esterase, thus preventing it from reaching receptors on SM cells. Consequently, we considered the enteric neural stimulus (*β*) to act exclusively on ICCs in our GMN model. The animal models that lack ICCs with muscarinic receptors show that there is little or no cholinergic response in SM cells because Acetylcholine is broken down before it can be taken up by receptors on SM cells [26]. In our model, we maintained a linear gradient of *β* to ensure that rostral ICCs successively entrained caudal ICCs. This way, we assembled a chain of realistic ICCs which show a rostrocaudal gradient in their intrinsic frequencies like that observed in the stomach. In the heart, fast-pacing sinoatrial node cells entrain the slow-pacing atrioventricular node cells [84], like what is observed in an intact stomach. Besides being intrinsic pacemakers, ICCs are also involved in neurotransmission, setting the membrane potential of SM cells, and in stretch sensing [51]. Future experiments examining effects of neural input on ICCs may shed further light on the neural contribution to pacemaker potential generation and slow-wave propagation. Empirical measurements of the neural stimulus on ICCs at different points along the stomach would strengthen the validity of our model, offering further hypotheses for the mechanisms underlying genesis of functional and pathological slow waves in the stomach. The mathematical modeling framework outlined in our study is a step in this direction and provides an exploration testbed for precise modulation of intra/intercellular pathways to examine their role in the above-mentioned phenomena.

## Methods

The model network of ICC-SM cells is described in **Fig 2**. The network consists of 42 ICCs and 42 SM cells implemented and simulated in MATLAB 2019a (Mathworks.inc). Together, the extensive biological details of ICC and SM cell physiology and the assumed intercellular coupling resulted in a large network model with 1554 nonlinear differential equations (23 for each ICC and 14 for each SM cell) consisting of over 1000 parameters. The parameter values were chosen from published models [7,42,43].

### ICC cell model

We adopted a well-described conductance-based model by Corrias *et al*., 2008 [42] for the ICCs (also see [7,19]). The rate of change in the membrane potential, *V*_*ICC*_, for each ICC is as follows:

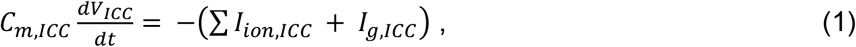

where *V*_*ICC*_ is the ICC membrane potential, *C*_*m,ICC*_ is the ICC membrane capacitance, *ICC*_*ion,ICC*_ represents the summation of all the ionic currents in ICC, and *ICC*_*g,ICC*_ represents the current through the gap junctions between the ICCs. The different ionic currents included in the model are described in **Table 1**. The dynamics of the voltage-dependent gating variables for the ionic currents follow the well-known Hodgkin-Huxley formalism, where each current is represented by a battery (electrochemical driving force) in series with a variable resistance and the cell membrane as a capacitor in parallel. The combined actions of these ion channel currents reproduce the three phases of a gastric action potential: depolarization, plateau phase, and repolarization. Detailed descriptions of these ionic currents are provided in the github repository (https://github.com/ashfaq-polit/Slow_waves_in_the_stomach) and are based on previous modeling studies [42,43,85].

### Ca^2+^-IP_3_ dynamics in ICCs

The equation guiding the intracellular Ca^2+^ dynamics is modeled similar to Fall and Keizer [86] as follows

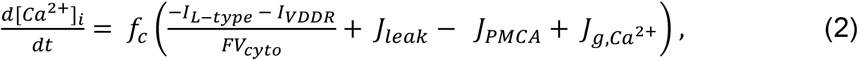

Where *I*_*L*−*type*_ and *I*_*VDDR*_ are described in Table 1, *J*_*leak*_ is a leakage flux between the pacemaker region and the cytosol, *J*_*PMCA*_ is the Ca^2+^ flux through the plasmalemmal Ca^2+^ pump, *F* is the Faraday constant, *f*_*c*_ is a dimensionless constant, *V*_*cyto*_ represents the cytosolic volume fraction within the ICC, and 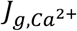 is the Ca^2+^ flux due to inter-ICC coupling (see **Equation 6**).

The Ca^2+^ flux in a sub-membrane space triggered by IP_3_ -operated stores in the endoplasmic reticulum (ER) is important for initiating a gastric action potential. The Ca^2+^ concentration in the sub-membrane space is modeled by the following equation:

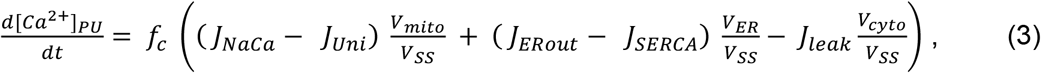

Where *J*_*N*a*Ca*_ and *J*_*Uni*_ represent mitochondrial Ca^2+^ fluxes, *J*_*ERout*_ and *J*_*SERCA*_ represent outward Ca^2+^ fluxes from the ER and inward Ca^2+^ fluxes through the ER. *V*_*mito*_, *V*_*ER*_, and *V*_*ss*_ represent the volume fractions for the mitochondria, endoplasmic reticulum, and pacemaker submembrane space, respectively.

Increases in intracellular IP_3_ concentration in each ICC are assumed to depend on: 1) voltage-dependent IP_3_ increase, 2) inter-ICC coupling-dependent IP_3_ increase, and 3) neurotransmitter-induced IP_3_ increase, whereas linear and non-linear degradation of IP_3_ decrease its concentration. Based on these assumptions, the rate of change in IP_3_ concentration is given as follows:

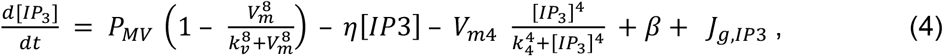

where *P*_*MV*_ is maximal rate of voltage-dependent IP_3_ synthesis, *K*_*V*_ is the half-saturation constant for voltage-dependent IP_3_ synthesis, *η* is the rate constant for linear IP_3_ degradation, *V*_*m*4_ is maximal value for the nonlinear IP_3_ degradation, *K*_4_ is half-saturation constant for the nonlinear IP_3_ degradation, *α* represents an enteric neural stimulus that modulates IP_3_ production, and *J*_*g,IP*3_ is the IP_3_ flux due to inter-ICC coupling (see **Equation 7**).

### Intercellular coupling between ICCs

Adjacent ICCs are assumed to be coupled via gap junctions with electrical conductance for passive ionic movement between cells [17]. The inter-ICC gap junction current is:

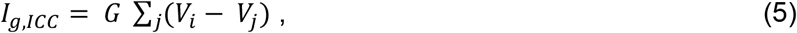

where *i* denotes the index for the source ICC and j denotes the index for adjacent ICCs; *G* represents electrical conductance of gap junctions.

### Second messenger exchange between ICCs

An exchange of second messengers, namely, Ca^2+^ and IP_3_ can occur via gap junctions [22– 24,81]. The flux describing Ca^2+^ exchange between adjacent ICCs is modeled by a term

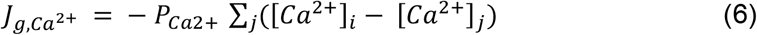

complementing Equation 2. Here *P*_*Ca*2+_ denotes the permeability coefficient for Ca^2+^ and [*Ca*^2+^]_*x*_ are the intracellular concentrations of Ca^2+^ in the corresponding cells where *x = i/j*. The 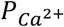 value has been taken from a theoretical study done in hepatocytes [87].

The inter-ICC IP_3_ flux is modeled using a similar formalism:

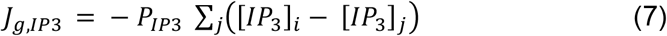

Here *P*_*IP*3_ denotes the permeability coefficient for IP_3_ and [*IP*_3_]_*x*_ are the intracellular concentrations of IP_3_ in the corresponding cells where *x = i/j*. An experimentally determined value of *P*_*IP*3_ is rare if not non-existent due to the technical difficulties of measuring [IP_3_] in tissue preparations. Hence, the permeability values used in model simulations were adjusted similar to [24,35].

### SM cell model

The SM cell model was adopted from Corrias *et al*., 2007 [43]. Like ICCs, the conductance-based rate of change in membrane potential is given as follows:

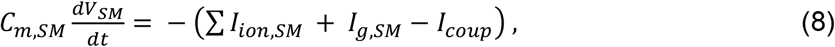

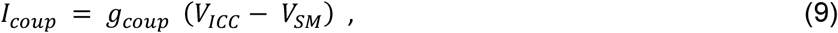

Where *V*_*SM*_ is the SM cell membrane potential, *C*_*m,SM*_ is the SM cell membrane capacitance, *I*_*ion,SM*_ represents the summation of all the ionic currents in the SM cell, *I*_*g,SM*_ represents the current through the gap junctions between the SM cells, and *I*_*coup*_ represents the coupling current from ICC to its corresponding SM cell. Although a bidirectional movement of ions can occur via gap junctions, here we assume that during entrainment depolarization spreads unidirectionally from ICC to SM and that *V*_*ICC*_ is always more positive than *V*_*SM*_ [70]; hence, the coupling current, *I*_*coup*_ is present only in **Equation 8**. The various ionic currents necessary to generate the SM action potential are listed in **Table 1**.

### Model simulation and analysis

The ICC-SM model network consisting of 42 ICCs and 42 SM cells was implemented and simulated in MATLAB 2019a (Mathworks.inc) using built-in function ODE15s and variable step-size. The total runtime for each simulation was 900 seconds (each simulation lasted approximately 8 hours on an Intel Xeon (R) CPU, 32 GB RAM Desktop computer). The outcome of the network was interpreted from the spatiotemporal membrane potential, *Relative Lag* and *period of SM cells* (explained in **Results** section). Statistical tests (t-test, one-way ANOVA) were performed as required and an *α* cutoff of 0.05 was chosen for statistical significance.

## Author Contributions

Conceptualization: RJ, MAA. Formal analysis: MAA. Investigation: MAA, SV, RJ. Project administration and Supervision: RJ. Visualization: MAA, SV. Writing of original draft: MAA, Review and Editing: MAA, SV, RJ.

## Supporting information

**S1 Table.**
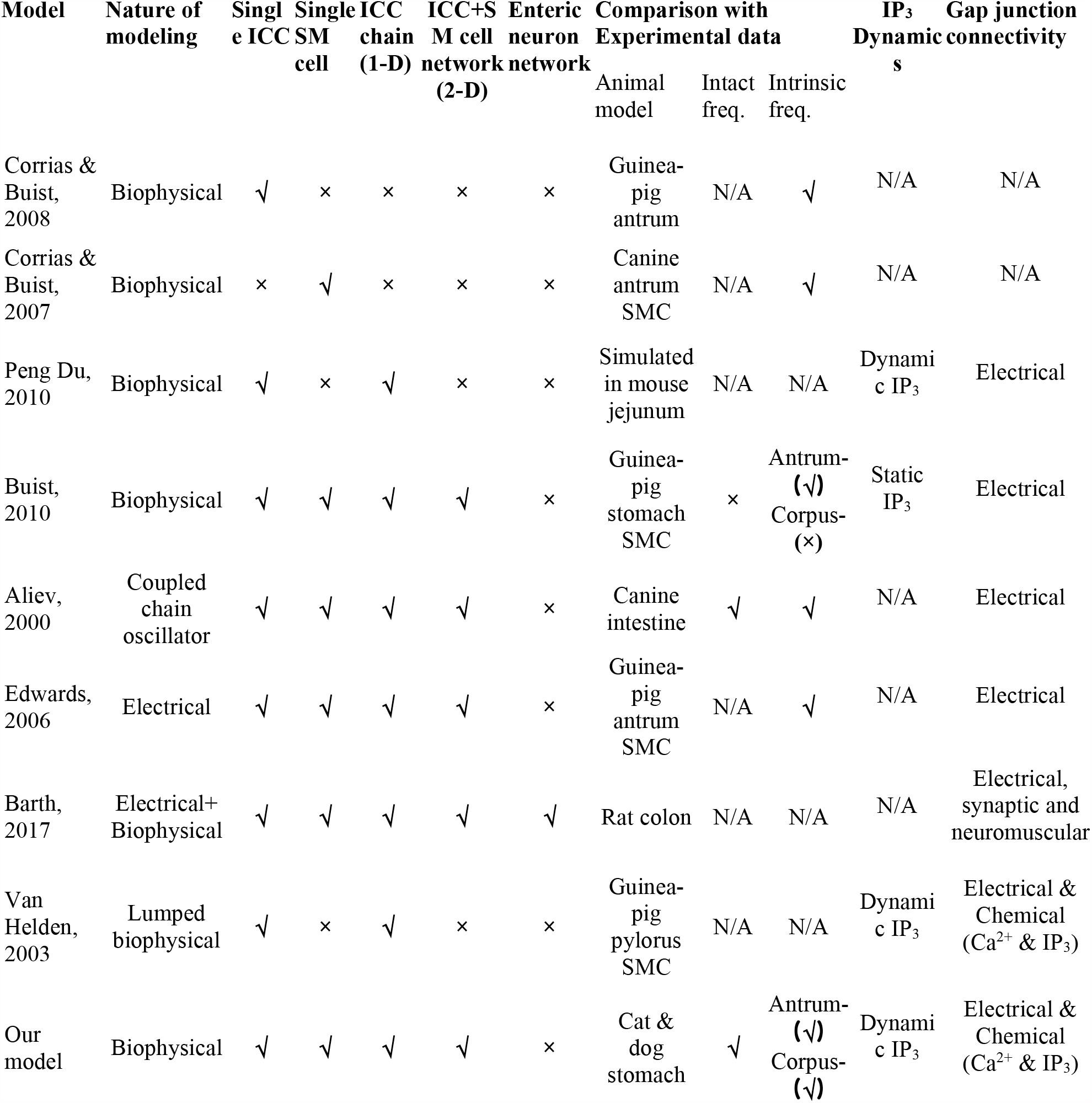
Summary of published models for a gastric slow-wave in the stomach.

**S2 Table.** *Total Lag* under different values of *G*_*ICC*−*ICC*_. Mean ± standard deviation for the last 7 cycles in each simulation.

| **$G_{ICC-ICC}$ value (nS)** | **Total Lag (sec)** |
| --- | --- |
| 0.35 | 53.77 ± 3.63 |
| 0.53 | 21.99 ± 0.02 |
| 0.7 | 20.85 ± 0.02 |
| 1.05 | 19.27 ± 0.03 |
| 1.4 | 18.22 ± 0.03 |
| 2.0 | 16.93 ± 0.03 |
| 3.0 | 15.51 ± 0.03 |
| 5.0 | 13.81 ± 0.03 |

**S3 Table.** *SM Cell Period* under different values of *G*_*ICC*−*ICC*_. Mean ± standard deviation for the last 7 cycles in each simulation.

| **$G_{ICC-ICC}$ value (nS)** | **SM<sub>1</sub> Period (sec)** | **SM<sub>42</sub> Period (sec)** |
| --- | --- | --- |
| 0.35 | 17.75 ± 0.01 | 19.41 ± 0.44 |
| 0.53 | 17.75 ± 0.01 | 17.74 ± 0.03 |
| 0.7 | 17.74 ± 0.01 | 17.73 ± 0.01 |
| 1.05 | 17.70 ± 0.01 | 17.69 ± 0.01 |
| 1.4 | 17.67 ± 0.01 | 17.66 ± 0.01 |
| 2.0 | 17.64 ± 0.01 | 17.62 ± 0.01 |
| 3.0 | 17.59 ± 0.01 | 17.58 ± 0.01 |
| 5.0 | 17.52 ± 0.01 | 17.51 ± 0.01 |

**S4 Table.** *Total Lag* under different values of *ICC*_3_. Mean ± standard deviation for the last 7 cycles in each simulation.

| $P_{IP3}$ value ($\text{sec}^{-1}$) | Total Lag (sec) |
| --- | --- |
| 4.0 | $69.56 \pm 2.45$ |
| 6.0 | $53.99 \pm 4.03$ |
| 8.0 | $20.85 \pm 0.02$ |
| 12.0 | $16.85 \pm 0.03$ |
| 16.0 | $13.17 \pm 0.03$ |
| 20.0 | $10.92 \pm 0.02$ |
| 30.0 | $8.90 \pm 0.01$ |
| 40.0 | $7.93 \pm 0.03$ |

**S5 Table.** *SM Cell Period* under different values of *P*_*IP*3_. Mean ± standard deviation for the last 7 cycles in each simulation.

| $p_{IP3}$ value ($\text{sec}^{-1}$) | $SM_1$ Period (sec) | $SM_{42}$ Period (sec) |
| --- | --- | --- |
| 4.0 | $17.31 \pm 0.01$ | $18.44 \pm 0.34$ |
| 6.0 | $17.53 \pm 0.02$ | $19.31 \pm 0.59$ |
| 8.0 | $17.74 \pm 0.01$ | $17.73 \pm 0.01$ |
| 12.0 | $17.94 \pm 0.01$ | $17.92 \pm 0.01$ |
| 16.0 | $18.31 \pm 0.01$ | $18.30 \pm 0.01$ |
| 20.0 | $18.58 \pm 0.01$ | $18.57 \pm 0.01$ |
| 30.0 | $18.69 \pm 0.02$ | $18.69 \pm 0.01$ |
| 40.0 | $18.71 \pm 0.03$ | $18.69 \pm 0.02$ |

**S6 Fig.**
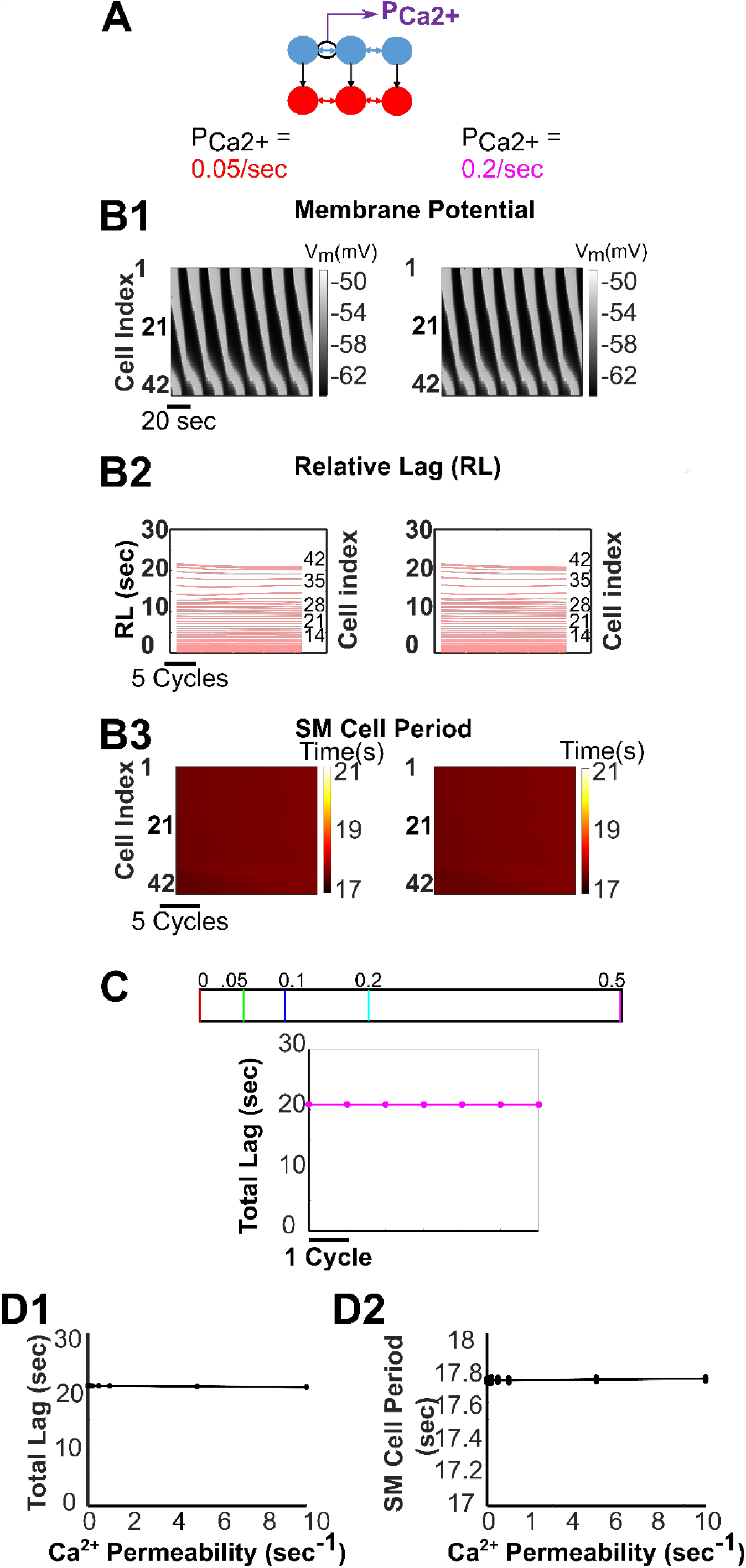
Increasing Ca^2+^ permeability across ICC-ICC gap junctions has no impact on network behavior. **(A)** The network. **(B)** Spatiotemporal map of membrane potential **(B1)**, *Relative Lags* **(B2)**, and spatiotemporal map of *SM Cell Periods* **(B3)** for the network when *P*_*Ca*2+_ = 0.05 sec^-1^ (left panel diagrams) and 0.2 sec^-1^ (right panel diagrams). **(C)** The *Total Lags* for changes in *P*_*Ca*2+_ are shown for the last 7 cycles of 900-sec simulations. The corresponding values of these permeabilities in sec^-1^ are shown in the legend. **(D)** For several networks, the mean *Total Lag* **(D1)** and the *SM Cell Period* **(D2)** of the last 7 cycles for each network with respect to its *P*_*Ca*2+_ are fit by an approximately constant line.

**S7 Fig.**
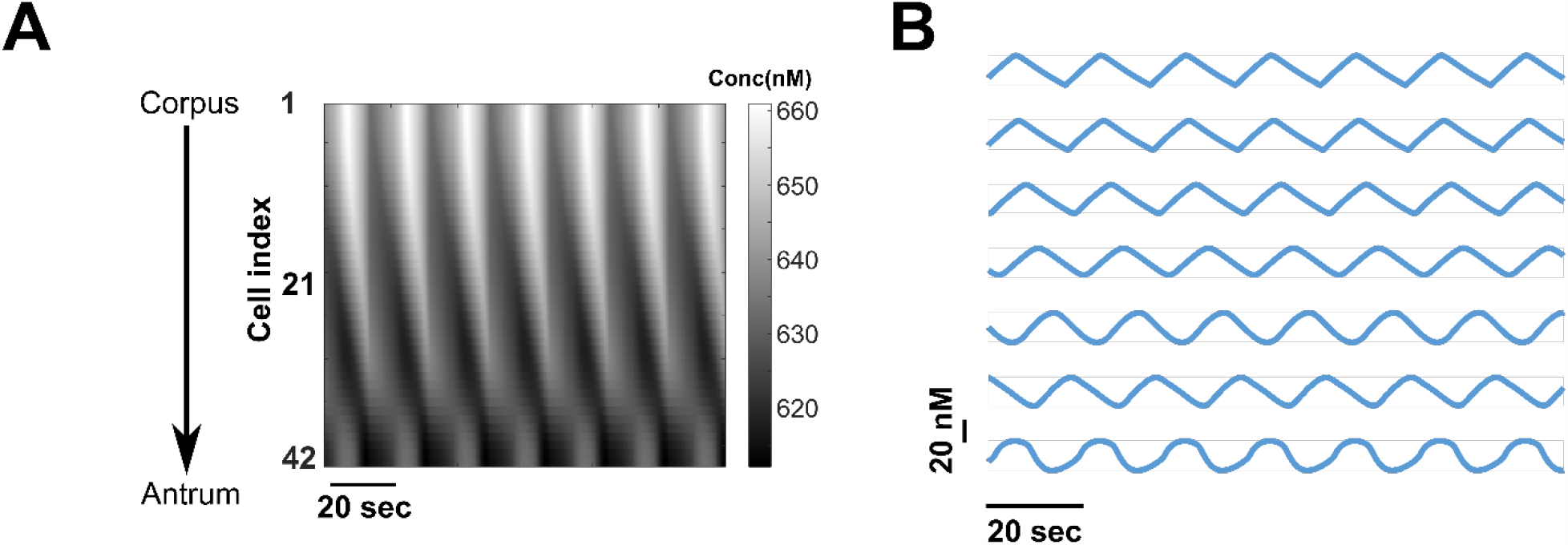
Spatiotemporal change in intracellular IP_3_ concentration. **(A)** Heatmap for all the cells in the network, and **(B)** for every 7^th^ cell in the network in steady-state.

**S8 Fig.**
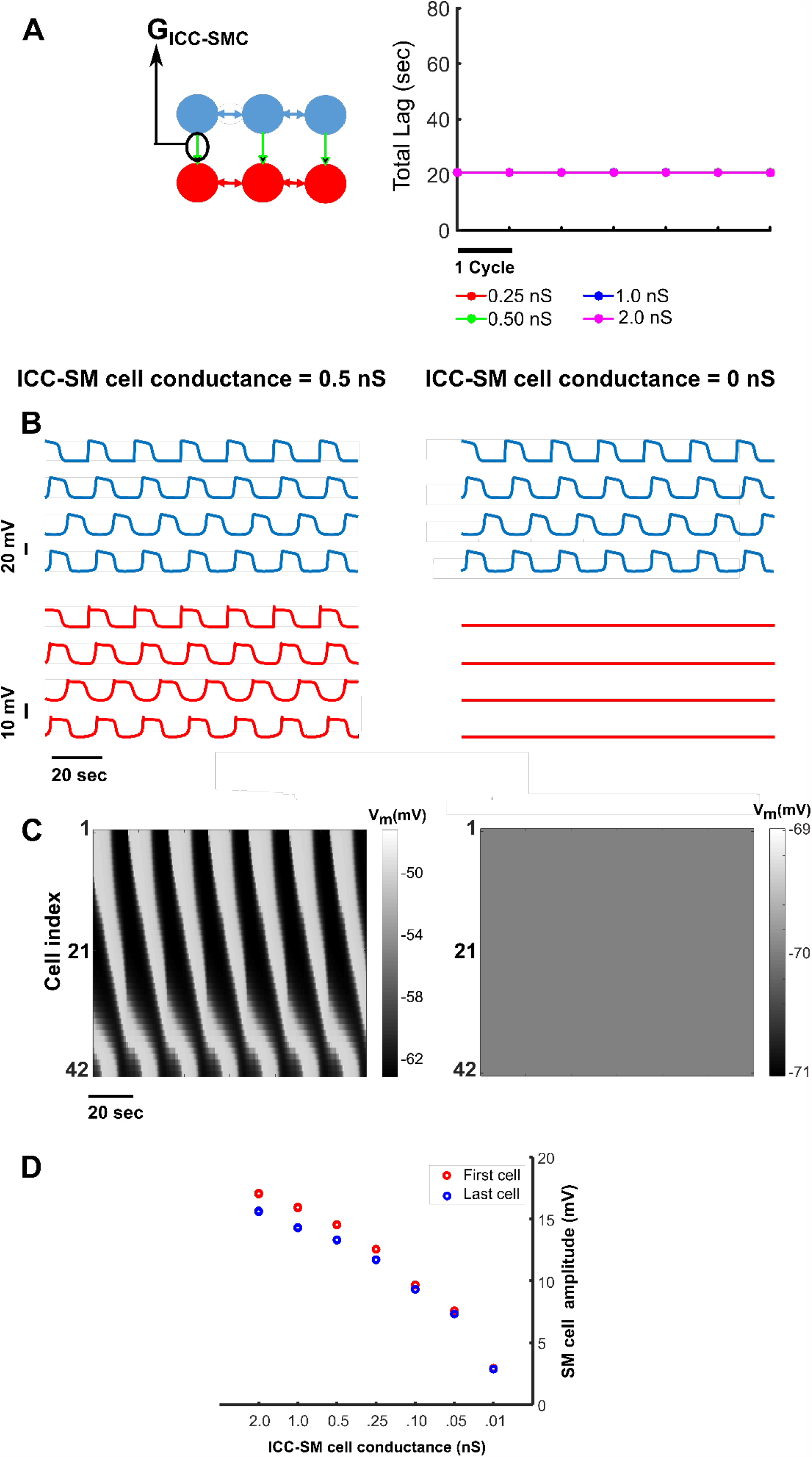
The ICC-SM cell gap junction conductance impacts the SM cell membrane potential amplitude. **(A)** ICC-SM cell electrical conductance does not have any effect on network entrainment evident from the approximately equal values of *Total Lag* measured for 4 different values of ICC-SM cell electrical conductances. **(B)** Membrane potential of 4 equidistant ICCs (top diagrams) and SM cells (bottom panel diagrams) in the 42-cell network when ICC-SM cell conductance is 0.5 ns (**left**) and 0 ns (**right**). **(C)** Spatiotemporal map of membrane potential of all 42 SM cells of the network, where ICC-SM cell conductance is 0.5 nS (**left**) and 0 nS (**right**). **(D)** Reduction of ICC to SM cell gap junction conductance reduces the amplitude of SM cell membrane potentials. For representation purposes, here the amplitudes (the peak-to-valley) of membrane potentials of the first and last SM cells of the network are shown.

